# Brassinosteroids signaling component SlBZR1 promotes fruit ripening in tomato

**DOI:** 10.1101/2022.07.11.499596

**Authors:** Fanliang Meng, Haoran Liu, Songshen Hu, Chengguo Jia, Min Zhang, Songwen Li, Yuanyuan Li, Jiayao Lin, Yue Jian, Mengyu Wang, Zhiyong Shao, Yuanyu Mao, Lihong Liu, Qiaomei Wang

**Affiliations:** Key Laboratory of Horticultural Plant Growth and Development, Ministry of Agriculture, Department of Horticulture, Zhejiang University, Hangzhou 310058, PR China; College of Plant Science, Jilin University, Changchun 130062, Jilin, PR China

**Keywords:** tomato, brassinosteroid, *SlBZR1*, fruit ripening, ethylene, carotenoids

## Abstract

Fruit ripening evolved to be attractive to frugivores that derive energy and nutrition from the fruits in exchange for assisting seed dispersal, which is accompanied by the dramatically change of fruit characteristics, including color, aroma, and texture. The plant hormone ethylene plays a key role in climacteric fruit ripening, while the role of other phytohormones as well as their cross talk with ethylene in modulating fruit ripening remains elusive. Here, we report growth-promoting phytohormone brassinosteroids promote fruit ripening in tomato through regulation of ethylene biosynthesis. Exogenous BR treatment and the increase of endogenous BR content in *SlCYP90B3*-OE promoted ethylene production and fruit ripening. SlBZR1, a central component and positive regulator of BR signaling pathway, promotes ethylene production and carotenoid accumulation through direct transcriptional regulation of *SlACO1, SlACO3* and *SlPSY1*. Furthermore, SlBIN2, a negative regulator of BR signaling upstream of SlBZR1, decreases ethylene production and carotenoid accumulation. Together, our results demonstrate that BR signaling integrates ethylene and carotenoid biosynthetic pathway to regulate fruit ripening.

## Introduction

Tomato (*Solanum lycopersicum*) is one of the most important vegetables as well as a perfect model system to study fruit ripening. Tomato fruit ripening is an intricate process in plant development that also involves overall fruit quality changes, such as color, texture, favor, and aroma [1]. In line with these changes, maturing tomato fruits undergo a series of physiological and metabolic processes, such as respiratory climacteric, cell wall synthesis and degradation as well as carotenoid biosynthesis [2]. Carotenoids are a diverse group of isoprenoid compounds with unique physicochemical properties, which are widely present in ripe tomato fruits, functioning as health-promoting natural pigments that participate in the coloration and quality formation of tomato fruits, as well as assisting seed dispersal by animals [3]. Significant progress has been achieved in recent years on carotenoid biosynthesis and catabolism pathway. *SlPSY1*, encoding phytoene synthase, is a key rate-limiting gene for carotenoid synthesis in tomato. Overexpression of *SlPSY1* accumulates elevated levels of carotenoids in tomato fruits [4]. Carotenoid metabolism in tomato fruits is closely synchronized with the ripening process, which is modulated by the interplay of various factors including developmental cues, environmental signals and phytohormones [3].

As a phytohormone, ethylene is an overall ripening-promoting regulator governing normal ripening and carotenoid accumulation of tomato fruits via a burst of ethylene production followed by an increasing in respiration rate at the onset of fruit ripening [1, 5, 6]. In higher plants, ethylene biosynthesis begins with the supply of S-adenosyl methionine (SAM) via the Yang pathway, which is catalyzed by ACC synthase (ACS) and converted into ACC, subsequent reaction catalyzed by ACC oxidase (ACO) eventually lead to the production of ethylene [7-9]. Previous studies manifest that the expression of *SlACO1* and *SlACO3* is markedly increased when tomato fruit ripening is triggered. Antisense-mediated RNA silencing of *SlACO1* causes delayed ripening and reduced biosynthesis of endogenous ethylene [10-12]. In addition, great progress has been made in identification of transcription factors responsible for tomato fruit ripening and carotenoid accumulation in an ethylene-dependent manner. Several tomato MADS-box transcription factors and NAC family transcription factors are reported to be involved in ethylene biosynthesis, while also affect the transcription of carotenoid biosynthetic genes. These tomato MADS-box transcription factors, including Ripening Inhibitor (RIN), Agamous-like1 (TAGL1), Fruitfull1 (FUL1), and FUL2 contribute to fruit ripening and carotenoid accumulation with the positive regulations on both ethylene biosynthesis genes, such as *ACS2* and *ACS4*, as well as carotenoid biosynthesis gene *SlPSY1* [13-15]. Moreover, RIN could also interact with TAGL1, FUL1, and FUL2 to form complex which functions as regulator of fruit ripening [13, 16-18]. NAC family transcription factors, including NOR and NOR-like1 positively modulate biosynthetic pathway of ethylene and carotenoid by directly binding to the promoter of *ACS2* and *SlGGPPS2* [19, 20]. Furthermore, other transcription factors such as CNR, AP2a, SlHB1, SlERF6, SGR1 as well as some WRKY transcription factors including SlWRKY35 and SlWRKY32 are also involved in ethylene biosynthesis and carotenoid accumulation during tomato fruit ripening [21-28]. The regulatory mechanism of ethylene in ripening of climacteric fruit, represented by tomato, has been well elucidated. However, fruit ripening is a complex developmental process fine-tuned by various factors, of which the interaction networks of phytohormone signaling play a central role, thus it will be interesting and significant to reveal the function of other phytohormones act either independently or in combination with ethylene in fruit ripening [24]. Moreover, the mechanism of ethylene promoting fruit ripening is primarily applied to extending the shelf life of fruits instead of regulating fruit quality till now. Therefore, elucidating the regulatory mechanism of phytohormone interaction network in fruit ripening will extend our understanding of fruit development as well as provide new strategies for fruit quality improvement.

To date, several phytohormones have been observed to influence ethylene biosynthesis and carotenoid accumulation during fruit ripening. Our previous studies found that Jasmonic acid (JA) and its volatile methyl ester, MeJA, participate in regulation of fruit ripening and carotenoid accumulation partially via an ethylene-independent pathway [29, 30]. Additionally, JA was also reported to promote ethylene production in apple fruit via direct transcriptional activation of both *MdACS1* and *MdACO1* by MdMYC2 during fruit ripening [31]. Abscisic acid (ABA) concentration gradually increased at the onset of tomato fruit ripening, and application of exogenous ABA promoted ethylene biosynthesis and fruit ripening [32]. Furthermore, treatment of auxin analogue 2,4-D can delay the ripening process of tomato fruit via inhibiting the production of ethylene and enhance the final contents of carotenoids in mature tomato fruits [33].

BRs are a class of steroid phytohormones with broad-spectrum of biological functions, and involved in diverse plant growth and developmental processes, such as seed germination, root growth, stem elongation, leaf morphogenesis, stomatal formation, floral development [34]. To date, the BR signaling pathway has been extensively explored by using multiple approaches. BRs are perceived at the plasma membrane by the extracellular domains of Brassinosteroid-Insensitive1 (BRI1) receptor and its co-receptor BRI1-associated receptor kinase 1 (BAK1). BRI1 and BAK1 trans-phosphorylate each other, which increases the activity of BRI1 resulting in BR signaling kinase 1 (BSK1) phosphorylation and its release from the receptor complex [35, 36]. Activated BSK1 binds to and activates BRI1-Suppressor 1 (BSU1), which leads to dephosphorylation and inhibition of the Brassinosteroid Insensitive 2 (BIN2) [37]. Subsequently, the downstream transcription factors such as Brassinazole Resistant 1 (BZR1) and BRI1-EMS Suppressor 1 (BES1) are dephosphorylated by a cooperative reaction of BIN2 and Protein Phosphatase 2A (PP2A), then accumulate in the nucleus and bind to the promoters of BR-responsive target genes, conferring a specific physiological response [38, 39]. When BRs are absent or present at low levels, BZR1 and BES1 are phosphorylated by active BIN2, which abolishes their DNA-binding activity and leads to their cytoplasmic retention by 14-3-3 proteins. When BRs are present at high levels, BZR1 and BES1 are dephosphorylated by protein phosphatase 2A (PP2A) and transported into the nucleus, then bind to the promoters of their target genes, resulting in gene activation or repression [40].

Recent studies have demonstrated that BRs exert effects on fruit development process, especially fruit maturation. The application of exogenous BR promotes the ripening of climacteric fruits, such as tomato, banana, persimmon and mango, as well as non-climacteric fruits, including strawberry and grape via different regulatory mechanisms [41-46]. Moreover, overexpression of the BR biosynthetic gene *DWARF, SlCYP90B3* and *GhDWF4* respectively in tomato, results in elevated level of endogenous BRs, increased ethylene production and carotenoid accumulation as well as earlier ripening in an ethylene-dependent manner [47-49]. Tomato fruits with enhanced BR signaling via over-expression of SlBRI1, the receptor gene, also displayed elevated ethylene production and carotenoid levels during fruit ripening [50]. These results indicate that BRs promotes tomato fruit ripening possibly through activating the expression of ethylene biosynthetic genes and ultimately the ethylene production. Additionally, our former research also showed that BR signaling promoted carotenoid accumulation and quality formation in tomato during fruit ripening period [51]. Several recent studies found that MaBZR1/2, PuBZR1 and DkBZR1 repress ethylene production, while DkBZR2 promotes ethylene production during postharvest fruit ripening [42, 52, 53].

Our previous reports demonstrated that exogenous application and increase of endogenous BR both promote fruit ripening and carotenoid accumulation, up-regulate the expression of ethylene and carotenoid biosynthetic genes, while the regulatory mechanism of BR signaling in biosynthesis of ethylene and carotenoid during tomato fruit ripening remains unclear. In the present study, with an aim to elucidate the role of BR signaling in tomato fruit ripening, we investigated the function of *SlBZR1* and *SlBES1*, the tomato closest homolog of *AtBZR1* and *AtBES1*. Interestingly, we found that double mutation via *SlBZR1*/*SlBES1*-KO delays tomato fruit ripening and ethylene release as well as reduces carotenoid accumulation. Overexpression of *SlBZR1* and knockout of *SlBIN2* promotes fruit ripening with elevated ethylene production and carotenoid levels. Further tests reveal that *SlBZR1* exerts direct transcriptional activation on ethylene biosynthetic genes *SlACO1* and *SlACO3* as well as carotenoid biosynthetic gene *SlPSY1* to promote ethylene and carotenoid biosynthesis and fruit ripening.

## Results

### BR promotes fruit ripening

Previous studies have shown that the accumulation of endogenous BRs increases during tomato fruit ripening, which indicates that BRs may have a positive effect on tomato fruit ripening [48]. To determine the role of BRs on tomato fruit ripening, we treated tomato fruits collected at mature green (MG) stage with 24-epibrassinolide (eBL) or propiconazole (PCZ) alone as well as the combination of eBL and PCZ (eBL + PCZ) at a concentration of 3 µM, respectively. Compared to the control fruits, eBL-treated fruits showed a significantly higher production of ethylene during storage. Similarly, the peak of climacteric ethylene production occurred about 1 day earlier than the control fruits, while the climacteric ethylene peak was significantly inhibited after PCZ treatment (Supplemental Figure S1A). Moreover, compared with the WT fruits, overexpression of BR biosynthetic gene *SlCYP90B3* increased the level of endogenous bioactive BR in fruits and accelerated fruit ripening, *SlCYP90B3-RNAi* transgenic lines delayed fruit ripening (Supplemental Figure S1B). In line with this, *SlCYP90B3-OE* advanced the onset of the peak of climacteric ethylene production by 1d, and the peak of ethylene production was higher than that in the WT fruits. In contrast, *SlCYP90B3-RNAi* caused a lower peak of ethylene production that occurred later about 2 days (Supplemental Figure S1C). The results demonstrate that exogenous eBL treatment and increase in endogenous bioactive BR levels exert a positive role in tomato fruit ripening.

### Fruit ripening was inhibited in tomato *bzr1bes1* double mutant

To better understand BR action and identify the candidate components of BR signaling pathway function in tomato fruit ripening, we manually searched for the *SlBZR1* and *SlBES1*, the homologous gene of *AtBZR1* and *AtBES1* in tomato, encoding proteins of 333 and 327 amino acids, with calculated molecular weights of 35.78 and 34.98 kDa, respectively. Amino acid alignment revealed that SlBZR1 and SlBES1 contain a DNA-binding domain with a nuclear localization signal sequence (NLS), a putative 14-3-3 binding site, and a putative PEST sequence, which are typical characteristics of BZR/BES transcription factors. Particularly, 25 serine/threonine residues recognized by the phosphorylation site of GSK-3 kinase BIN2 were also found in SlBZR1 and SlBES1 proteins. In addition, the EAR (ethylene-responsive element binding factor-associated amphiphilic repression) domain motif was found to be localized near the C terminus of SlBZR1 and SlBES1 proteins (Supplemental Figure S2). Phylogenetic tree based on BZR sequences indicated that SlBZR1 and SlBES1 closely clustered with Arabidopsis AtBZR1 and AtBES1 (Supplemental Figure S2).

To verify whether BRs promotes the ripening of tomato fruits via *SlBZR1* and *SlBES1*, we generated loss-of-function mutants of both *SlBZR1* and *SlBES1* using CRISPR/Cas9 system in tomato cultivar *Ailsa Craig* (AC) background. To this end, two specific target sites were designed in the second exon of *SlBZR1* and *SlBES1*. Genomic DNA sequencing revealed two null double mutants *bzr1bes1-1* and *bzr1bes1-2* carried long deletions and single-base deletions causing frameshift and premature termination of translation, likely resulting in complete loss-of-function. Then, two distinct homozygous mutants *bzr1bes1-1* and *bzr1bes1-2* were selected for further analysis, the editing details of the two lines are shown in Figure 1A.

**Figure 1.**
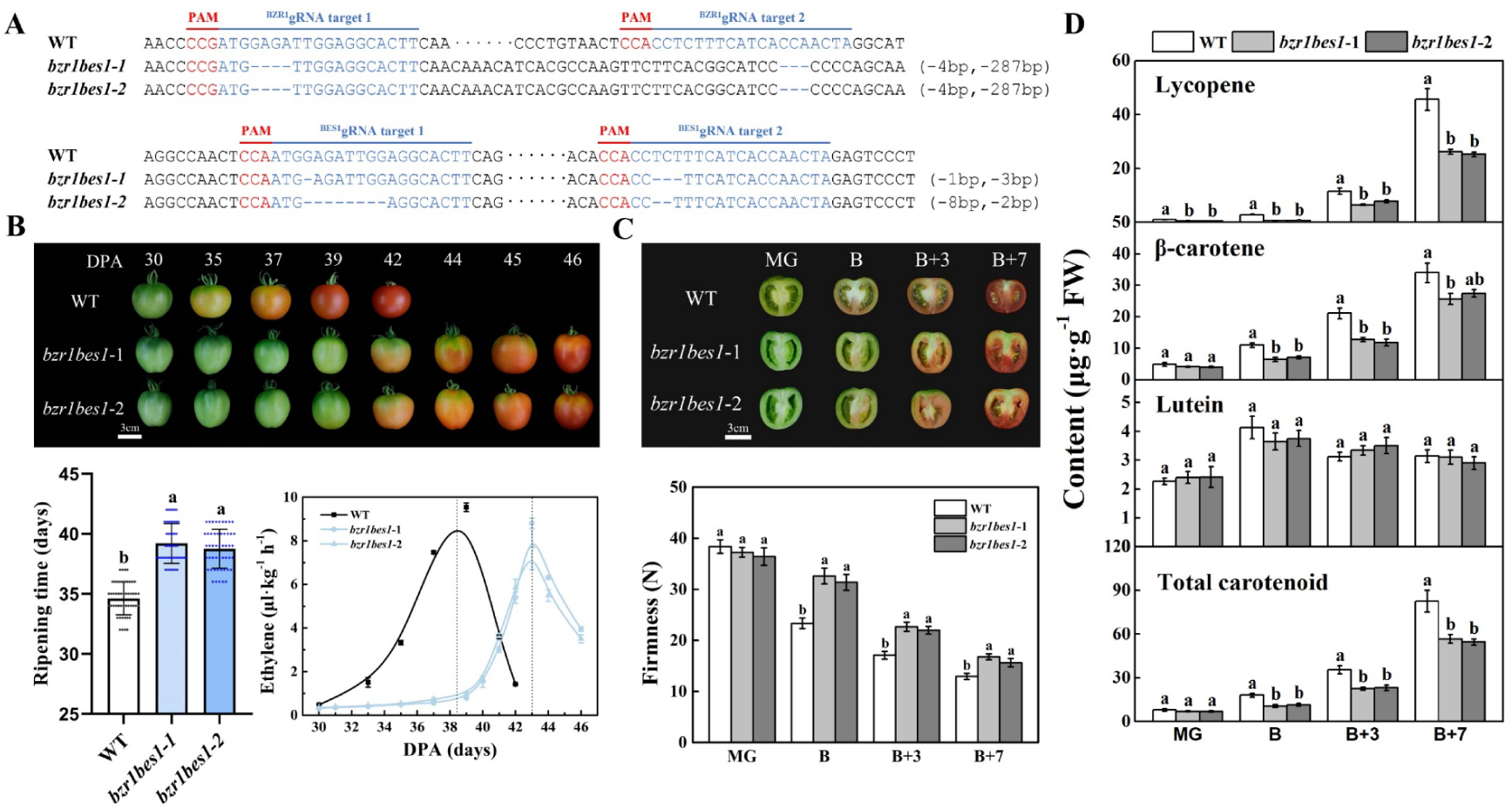
SlBZR1/SlBES1 double gene knockout abolish BR signaling delays tomato fruit ripening and ethylene production, reduces carotenoid accumulation. (A) Genotype of mutations in the SlBZR1 and SlBES1 locus generated by the CRISPR/Cas9 genome editing system. Two target sequences were designed to specifically target the second exons with a 221-bp interval. The red and blue letters indicate the sequences of protospacer adjacent motif (PAM) and target, respectively. The mutations in the transgenic plants were confirmed by sequencing the genomic regions flanking the target sites. The sequences of the wild-type (WT) plants and two homozygous mutant lines (*bzr1bes1-1* and *bzr1bes1-2*) are shown. (B) Ripening phenotype of *bzr1bes1* null mutant fruits. WT and *bzr1bes1* mutant fruits at 30, 35, 37, 39, 42, 44, 45, and 46 days post anthesis (DPA) are shown. Ripening time (days from anthesis to breaker stage) of WT and *bzr1bes1* mutant fruits. Changes in ethylene production of WT and *bzr1bes1* mutant fruits. (C) Longitudinal section phenotypes of WT and *bzr1bes1* mutant fruits at four different developmental stages. Fruit firmness and carotenoids content (D) were measured. MG, mature green stage; B, breaker; B + 3, 3 d after breaker stage; B + 7, 7 d after breaker stage. Biological replicates (3-4 fruits per fruit ripening stage) were performed in triplicate, and the data are presented as means ± SE. The asterisks indicate statistically significant differences between the WT and transgenic fruits (P < 0.05).

We observed and recorded the ripening characteristics of *bzr1bes1* mutant fruits. Notably, the fruit ripening progress was markedly delayed in *bzr1bes1-1* and *bzr1bes1-2* mutant lines compared with the WT. Fruits of both *bzr1bes1* double mutant lines required an additional 4-5 days from anthesis to the breaker stage compared with the WT fruits. Indeed, fruits of WT plants reached the breaker stage at 35-day post-anthesis (DPA), whereas the average time from anthesis to breaker stage extended to 40 DPA and 39 DPA in fruits of *bzr1bes1-1* and *bzr1bes1-2* lines, respectively. We next investigated climacteric ethylene production in *bzr1bes1* fruits by monitoring ethylene production from 30 to 46 DPA. The peaks of climacteric ethylene production in fruits of *bzr1bes1-1* and *bzr1bes1-2* lines appeared about 5 days later than in fruits of WT plants (Figure 1B).

To determine whether fruit quality was altered in *bzr1bes1* mutants, several typical fruit quality parameters were measured at different ripening stages (MG, B, B+3, and B+7). Fruits of both *bzr1bes1-1* and *bzr1bes1-2* lines exhibited increased fruit firmness compared with that of WT at B, B+3 and B+7 stages (Figure 1C), consistent with the delayed ripening of *bzr1bes1* fruits. The contents of lycopene, β-carotene and total carotenoids in fruits of *bzr1bes1-1* and *bzr1bes1-2* lines were much lower than those in fruits of WT at B, B+3 and B+7 stages (Figure 1D). These results suggest that SlBZR1 and SlBES1 play a positive role in fruit ripening and carotenoids accumulation.

### *SlBZR1* promotes tomato fruit ripening

Previous studies have shown that *SlBES1* promotes tomato fruit softening without affecting carotenoids contents and ethylene production (Supplemental Figure S3, A and B) [54], which indicates that *SlBZR1* might play a dominant role in tomato fruit ripening and carotenoids accumulation.

To gain insight into the biological function of SlBZR1 in tomato fruit ripening, we generated *SlBZR1*-overexpression transgenic lines and *SlBZR1* knock-out mutants using CRISPR/Cas9 system in AC. Three transgenic lines (OE-8, OE-9 and OE-14) with elevated *SlBZR1* gene expression levels were selected for further phenotypic and molecular analyses (Supplemental Figure S4, A and B). Likewise, three homozygous *bzr1* mutants (*bzr1-5, bzr1-28* and *bzr1-29*) with deletions or insertions that led to translational frame shifts were obtained, which encode truncated SlBZR1 proteins with 58, 48 and 49 aa, respectively (Supplemental Figure S4C). These results suggest that *bzr1-5, slbzr1-28* and *slbzr1-29* are loss-of-function mutants suitable for functional analysis of *SlBZR1*.

The fruit ripening progress was increasingly promoted in *SlBZR1*-OE lines and inhibited in *bzr1* mutant lines when compared with the WT plants (Figure 2, A and B). Notably, all three *SlBZR1*-OE tomato lines displayed a significantly advanced onset of fruit ripening by 2-3 days, while knockout of *SlBZR1* delayed onset of fruit ripening by 1 day (Figure 2, B and C), consistent with the hypothesis that *SlBZR1* positively regulates tomato fruit ripening. The peaks of climacteric ethylene production occurred 1-3 days earlier in *SlBZR1*-OE fruits and 1-2 days later in *bzr1* mutant fruits than in the WT (Figure 2D).

**Figure 2.**
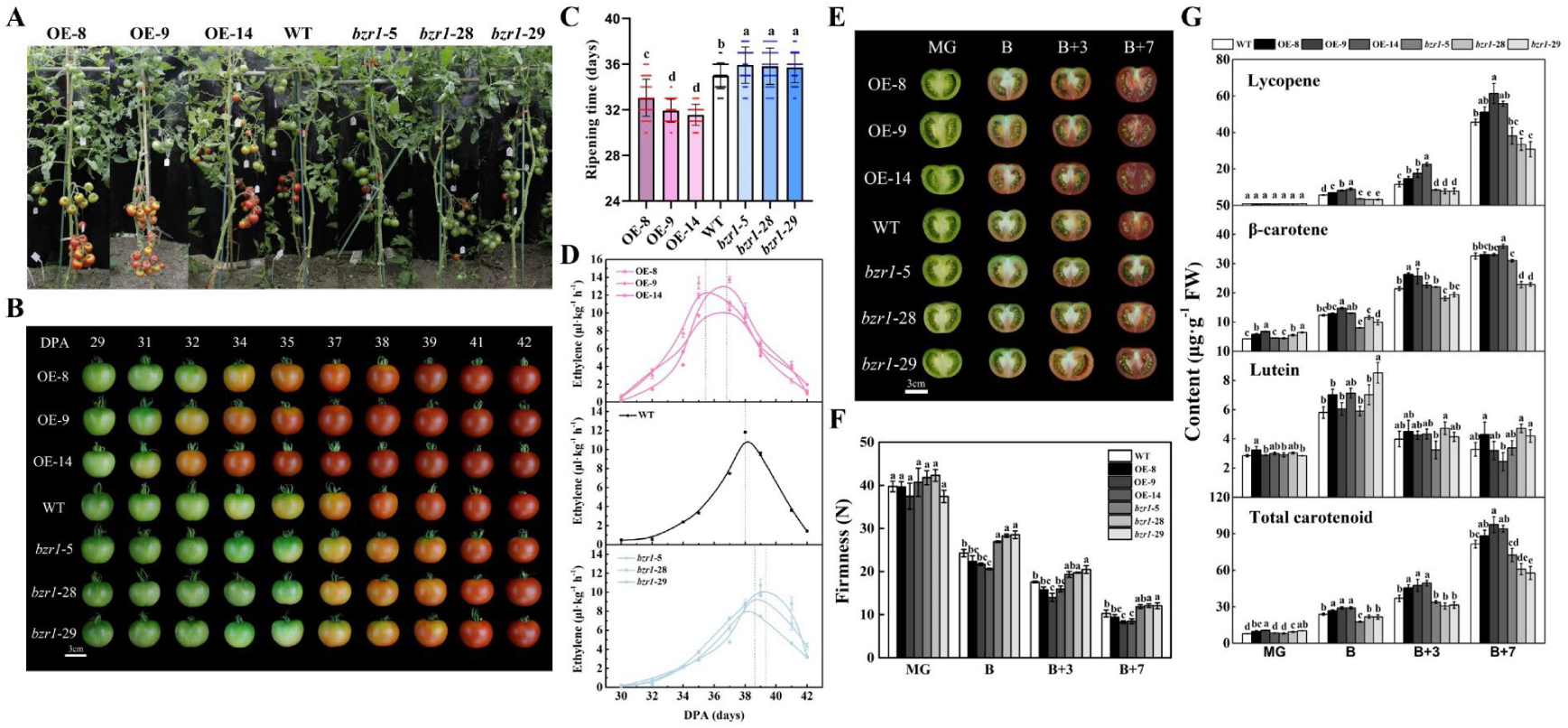
*SlBZR1* promotes the fruit ripening schedule and carotenoids accumulation. (A) Phenotypes of *SlBZR1*-OE and *bzr1* mutant lines. (B) Ripening phenotype of *SlBZR1*-OE and *bzr1* mutant fruits. Fruits from *SlBZR1*-OE lines ripen earlier while *bzr1* mutant lines show a delayed ripening phenotype. (C) Ripening time (days from anthesis to breaker) of WT and *SlBZR1*-OE and *bzr1* mutant fruits. (D) Ethylene production in WT, *SlBZR1*-OE and *slbzr1* mutant fruits at different developmental stages. (E) Cross-sectioned WT, *SlBZR1*-OE and *slbzr1* mutant fruits at four different developmental stages. (F) Fruit firmness and carotenoids content (G) in WT, *SlBZR1*-OE and *slbzr1* mutant fruits at four different developmental stages. MG, mature green stage; B, breaker; B + 3, 3 d after breaker stage; B + 7, 7 d after breaker stage. Biological replicates (3–4 fruits per fruit ripening stage) were performed in triplicate, and the data are presented as means ± SE. The asterisks indicate statistically significant differences between the WT and transgenic fruits (*P < 0.05).

To explore the effects of *SlBZR1* on major physiological characteristics of tomato fruit ripening, we also measured fruit firmness and carotenoid content of *SlBZR1*-OE, *bzr1* mutant and WT fruits. The results showed that the fruit firmness of *SlBZR1*-OE fruits was significantly reduced compared with that of WT fruits at B, B+3 and B+7 stages, while the fruit firmness of *bzr1* mutant fruits was enhanced than that of WT fruit at the same stage (Figure 2, E and F). The contents of lycopene, β-carotene and total carotenoids in *SlBZR1*-OE fruits were significantly higher than those in WT fruits, whereas *bzr1* mutant fruits produced less carotenoids than WT fruits, especially lycopene (Figure 2G). These results indicated that *SlBZR1* was a positive regulator of tomato fruit ripening and carotenoids accumulation.

### *SlBZR1* is a master regulator of tomato fruit ripening

To further elucidate the molecular mechanism of *SlBZR1* in regulating tomato fruit ripening and carotenoids biosynthesis, we performed global transcriptomic profiling by RNA-seq coupled with genome-wide ChIP-seq to identify genes and pathways potentially involved in tomato fruit ripening. *SlBZR1* knockout led to substantial transcriptomic reprogramming with 1,995 and 1,677 genes being differentially expressed (DEGs) in *bzr1* mutant fruits at MG and B stages, respectively. Among these, 730 genes were up-regulated and 1265 genes were down-regulated in *bzr1* mutant fruits at MG stage, 603 genes were up-regulated and 1074 genes were down-regulated in *bzr1* mutant fruits at B stage (Figure 3A). The up-regulated and down-regulated genes in *bzr1* mutant were termed SlBZR1-repressed genes and SlBZR1-induced genes, respectively.

**Figure 3.**
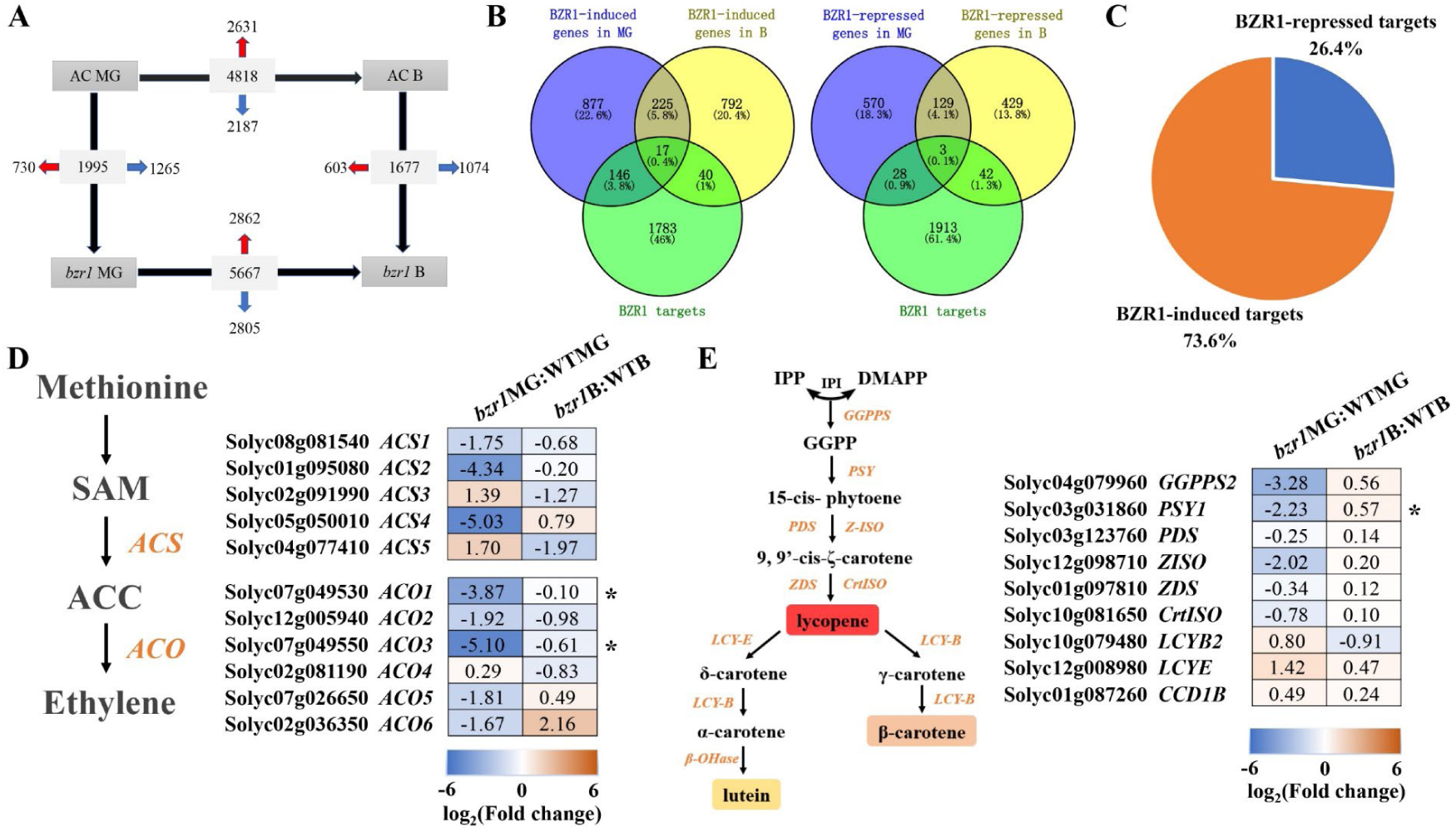
*SlBZR1* target genes identified by combined genome-wide RNA-seq and ChIP-seq approaches. (A) Number of differentially expressed genes (|log_2_(fold change)|>1 and p-value<0.05) between WT and *bzr1* at MG and B stage identified by RNA-seq. Indicated are total numbers (boxed) and numbers of up-regulated 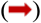 and down-regulated genes 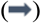 between stages and genotypes. (B) Venn diagram showing the numbers of genes common to *SlBZR1*-regulated genes and SlBZR1 target genes. (C) Percentage of SlBZR1-repressed and SlBZR1-induced target genes (in total 276). (D) Heatmap showing the differentially expressed genes involved in ethylene biosynthesis identified from the comparison of the *bzr1* mutant versus WT fruits at MG and B stage based on RNA-seq analysis. (E) Heatmap showing the differentially expressed genes involved in carotenoid biosynthesis identified from the comparison of the *bzr1* mutant versus WT fruits at MG and B stage based on RNA-seq analysis.

Moreover, 1986 potential SlBZR1 targets genes were identified by ChIP-seq using a polyclonal anti-GFP antibody and *SlBZR1*-OE fruits at MG stage expressing the BZR1-eGFP fusion protein. We compared the SlBZR1-repressed and SlBZR1-induced differentially expressed genes set obtained by RNA-seq with the SlBZR1 target genes set revealed by ChIP-seq. Cross-referencing the ChIP-seq and RNA-seq data identified 276 genes being both *SlBZR1* target and displaying differential expression in *bzr1* mutant fruits (Figure 3B). Interestingly, 73 genes (26.4%) were *SlBZR1*-repressed targets while 203 genes (73.6%) were *SlBZR1*-induced targets, consistent with *SlBZR1* functions as a positive regulator in tomato fruit ripening (Figure 3, B and C).

The production of ethylene and accumulation of carotenoids as well as fruit softening appears to be a typical feature during tomato fruit ripening. The RNA-seq and ChIP-seq analysis showed that numerous genes involved in fruit ripening including ethylene biosynthesis and carotenoid metabolism as well as transcription factor genes controlling fruit softening and fruit ripening were down-regulated in *bzr1* mutant fruits (Figure 3, D and E).

The fruit softening-related genes were *SlBZR1*-induced target genes and down-regulated in the *bzr1* mutant fruits, which indicates *SlBZR1* promoted tomato fruit softening by directly regulating fruit softening-related genes. Previous studies have demonstrated that SlBES1 promotes tomato fruit softening through transcriptional inhibition of *PMEU1* without affecting nutritional quality, which indicates that *SlBZR1* and *SlBES1* might perform partially redundant functions in fruit softening and regulate fruit ripening in a reciprocal manner.

The ethylene biosynthesis genes *SlACO1* and *SlACO3* as well as carotenoid biosynthesis gene *SlPSY1* were also *SlBZR1*-induced target genes and down-regulated in *bzr1* mutant fruits, indicating that *SlBZR1* promoted ethylene production and carotenoids accumulation by directly regulating these candidate ethylene and carotenoid biosynthesis genes (Figure 3, D and E). Moreover, the fruit ripening-related TFs were also *SlBZR1*-induced target genes and down-regulated in *bzr1* mutant fruits. These results were in agreement with the advanced onset of ripening in *SlBZR1*-OE fruits and the delayed ripening initiation in *bzr1* mutant fruits.

### Ethylene biosynthetic genes *SlACO1* and *SlACO3* as key targets of *SlBZR1* action

The expression levels of *SlACO1* and *SlACO3* were markedly increased in *SlBZR1*-OE and decreased in *bzr1* mutant fruits compared with the WT fruits at MG, B and P stages, consistent with the advanced onset of the peak of climacteric ethylene production (Figure 4, A and B). To understand whether SlBZR1 directly regulates transcription of *SlACO1* and *SlACO3* during tomato fruit ripening, we performed promoter analysis of *SlACO1* and *SlACO3* and found that there was one E-box and eight E-box in six regions in the 3-kb upstream regions of the promoters of *SlACO1* and *SlACO3*, respectively. Chromatin immunoprecipitation quantitative PCR (ChIP-qPCR) analysis showed a significantly higher enrichment for the promoters of *SlACO1* and *SlACO3*, which indicated that SlBZR1 directly bound to the promoters of both ethylene biosynthesis genes (Figure 4, C and D).

**Figure 4.**
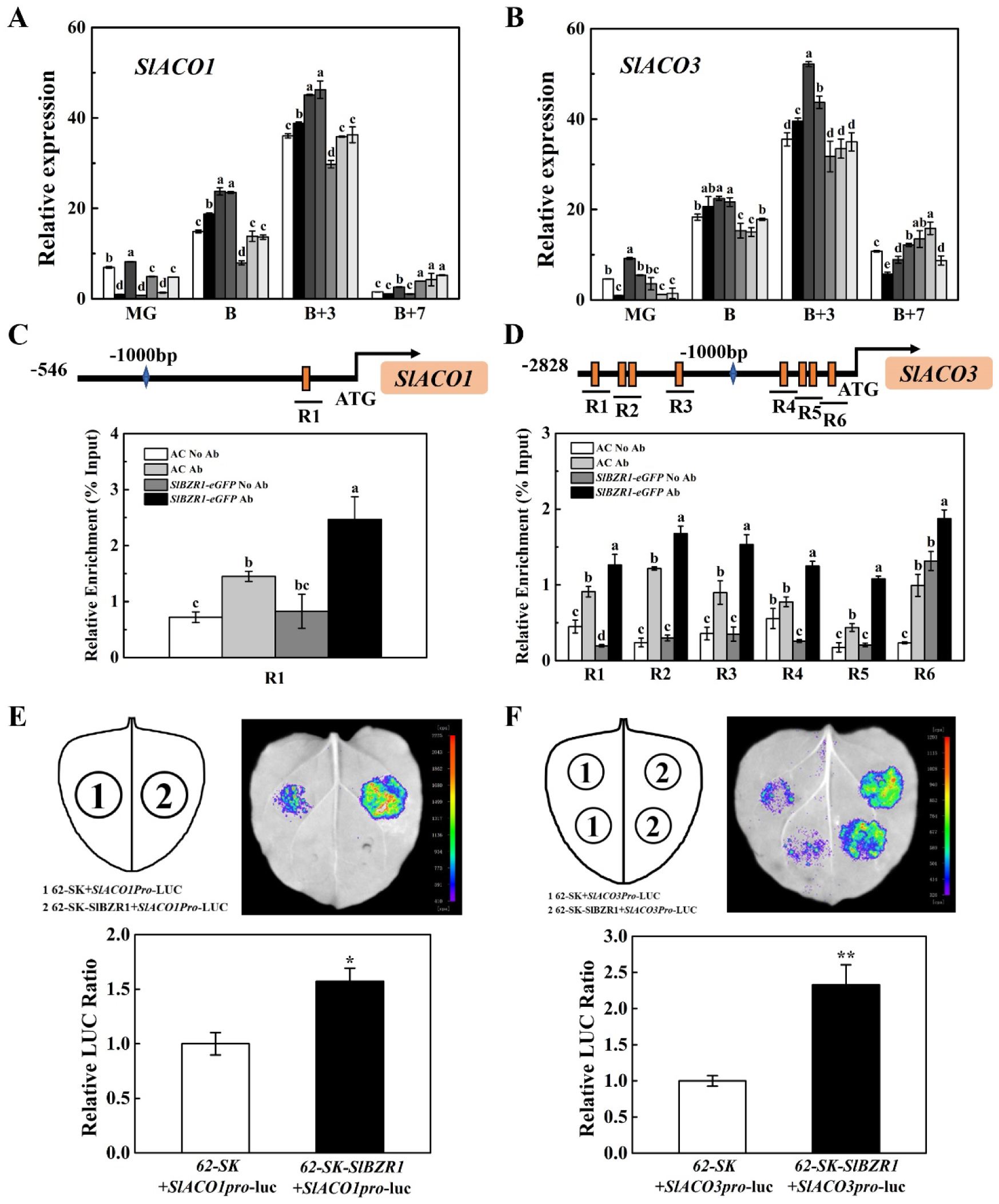
SlBZR1 directly binds and activates the expression of *SlACO1* and *SlACO3*. (A) Relative expression levels of *SlACO1* and *SlACO3* (B) in SlBZR1-OE and *bzr1* mutant fruits at different development stages. MG, mature green stage; B, breaker; B + 3, 3 d after breaker stage; B + 7, 7 d after breaker stage. Biological replicates (3-4 fruits per fruit ripening stage) were performed in triplicate, and the data are presented as means ± SE. The asterisks indicate statistically significant differences between the WT and transgenic fruits (P < 0.05). (C) ChIP-qPCR assays showing that SlBZR1 binds to the promoters of *SlACO1* and *SlACO3* (D). The promoter structures of the SlBZR1 target genes *SlACO1* and *SlACO3* are shown. The orange boxes represent E-box elements, and the numbers indicate the positions of these motifs relative to the translational start site. The black lines with uppercase letters represent the regions used for ChIP-qPCR. The relative enrichment values show the percentages of DNA fragments that coimmunoprecipitated with anti-GFP antibodies relative to the input DNAs. AC (wild type) without and with antibody were set as blank control and negative control, respectively. SlBZR1-GFP without antibody was also set as negative control. The mean value of two technical replicates was recorded for each biological replicate. Values are means ± SD of three biological replicates. Different letters indicate significant difference among groups for each locus (one-way ANOVA with Tukey’s test, P < 0.05). (E, F) Transient expression assay for SlBZR1 activation of the promoters of *SlACO1* and *SlACO3*. The reporter and effector vectors in each experiment were co-introduced into tobacco (*N. benthamiana*) leaves by Agrobacterium GV3101. SK represents empty vector control. Representative images of tobacco leaves were taken 3d after infiltration. Luminescence intensity values are means ± SD of six biological replicates. Different letters indicate significant difference among groups for each locus (one-way ANOVA with Tukey’s test, P < 0.05).

Next, we tested whether SlBZR1 could activate the transcription of *SlACO1* and *SlACO3* using the dual luciferase reporter assays. The promoter sequences of *SlACO1* and *SlACO3* were fused successfully to the LUC reporter gene, co-transformed with an empty pGreenII 62-SK vector or pGreenII 62-SK-SlBZR1 vector into Nicotiana benthamiana leaf epidermal cells. The results showed that SlBZR1 significantly activated the promoters of the *SlACO1* (1.57-fold) and *SlACO3* (2.33-fold) compared with the activity obtained with the control samples, respectively (Figure 4, E and F). Taken together, the results demonstrated that SlBZR1 positively regulates ethylene biosynthesis in tomato fruit by directly targeting the promoter of *SlACO1* and *SlACO3*.

### SlBZR1 promotes carotenoid accumulation in a partially ethylene-dependent manner

To further investigate the influence of the interaction between SlBZR1 and ethylene biosynthesis genes on promoting carotenoid accumulation during tomato fruit ripening, we treated *SlBZR1*-OE (OE-14) and WT fruits collected at mature green stage with 1 μL·L^-1^ 1-MCP for 12 h. The contents of lycopene and total carotenoids in *SlBZR1*-OE fruits were much higher than in WT fruits after 12 d and 18 d of storage, whereas the accumulation of lycopene, β-carotene and total carotenoids was significantly inhibited by 1-MCP treatment at 6 d, 12 d and 18 d of storage in both *SlBZR1*-OE and WT fruits. Interestingly, the promotion of lycopene and total carotenoid accumulation in the *SlBZR1*-OE fruits were not completely inhibited by 1-MCP treatment. The contents of lycopene and total carotenoids increased in the *SlBZR1*-OE fruits stored for 18 d after 1-MCP treatment compared with those in the wild-type fruits (Supplemental Figure S5, A and B). These results implied that *SlBZR1*-OE might promote carotenoid accumulation during tomato fruits ripening in a partially ethylene-dependent manner.

### SlBZR1 transactivates the expression of *SlPSY1* to regulate carotenoid accumulation

The expression levels of *SlPSY1* were markedly increased in *SlBZR1*-OE and decreased in *bzr1* mutant fruits compared with WT fruits at MG, B and P stages, which implied SlBZR1 might positively regulate the expression of *SlPSY1* (Figure 5A). To verify the direct regulation of BZR1 on *SlPSY1* expression, we performed promoter analysis of *SlPSY1* gene and found that there was three E-box in the 2-kb upstream regions of the promoters of *SlPSY1*. ChIP-qPCR analysis showed a significantly higher enrichment for the promoters of *SlPSY1*, which indicated that SlBZR1 directly bound to the promoters of *SlPSY1* (Figure 5B).

**Figure 5.**
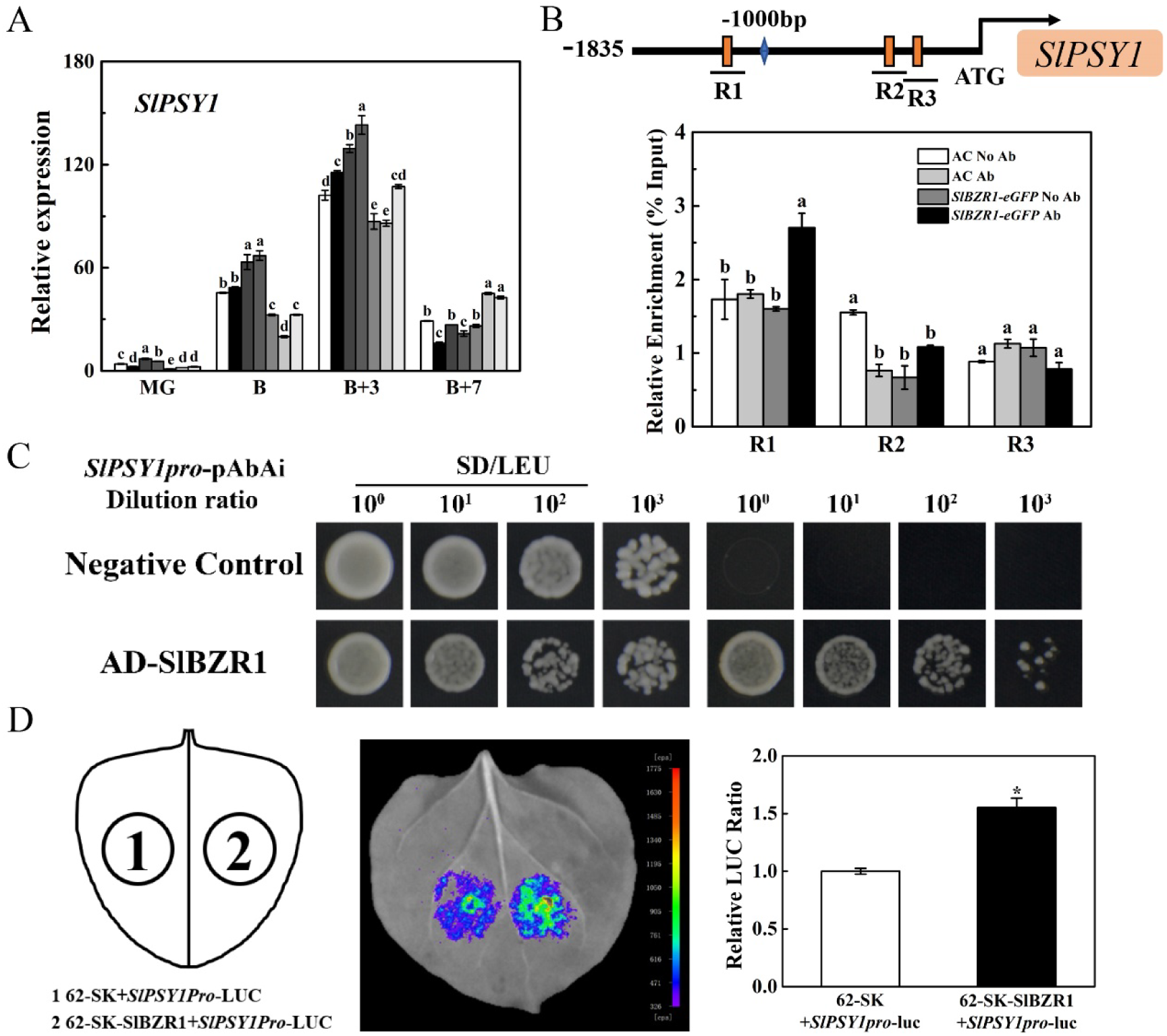
SlBZR1 positively regulates the expression of *SlPSY1* involved in carotenoid accumulation in ripening tomato fruits. (A) Relative expression levels of *SlPSY1* in SlBZR1-OE and *bzr1* mutant fruits at different development stages. MG, mature green stage; B, breaker; B + 3, 3 d after breaker stage; B + 7, 7 d after breaker stage. Biological replicates (3-4 fruits per fruit ripening stage) were performed in triplicate, and the data are presented as means ± SE. The asterisks indicate statistically significant differences between the WT and transgenic fruits (P < 0.05). (B) ChIP-qPCR assays showing that SlBZR1 binds to the promoters of *SlPSY1*. The promoter structures of the SlBZR1 target genes *SlPSY1* are shown. The orange boxes represent E-box elements, and the numbers indicate the positions of these motifs relative to the translational start site. The black lines with uppercase letters represent the regions used for ChIP-qPCR. The relative enrichment values show the percentages of DNA fragments that coimmunoprecipitated with anti-GFP antibodies relative to the input DNAs. AC (wild type) without and with antibody were set as blank control and negative control, respectively. SlBZR1-GFP without antibody was also set as negative control. The mean value of two technical replicates was recorded for each biological replicate. Values are means ± SD of three biological replicates. Different letters indicate significant difference among groups for each locus (one-way ANOVA with Tukey’s test, p < 0.05). (C) Y1H analysis showing that SlBZR1 binds to the promoter of *SlPSY1*. AbA, a yeast cell growth inhibitor, was used as a screening marker. The basal concentration of AbA was 150 ng·mL^−1^. The empty vector and the *SlPSY1* promoter were used as negative controls. (D) Transient expression assay for SlBZR1 activation of the promoters of *SlPSY1*. The reporter and effector vectors in each experiment were co-introduced into tobacco (N. benthamiana) leaves by Agrobacterium GV3101. SK represents empty vector control. Representative images of N. benthamiana leaves were taken 3d after infiltration. Luminescence intensity values are means ± SD of six biological replicates. Different letters indicate significant difference among groups for each locus (one-way ANOVA with Tukey’s test, p < 0.05).

Next, a yeast one-hybrid analysis was used to test for interaction between SlBZR1 and the promoter of *SlPSY1*. Yeast cells that were co-transformed with pGADT7-SlBZR1 and *SlPSY1pro-*pAbAi were able to grow on the SD/-Leu medium supplemented with AbA (150 ng·ml^-1^), whereas the negative control co-transformed with pGADT7 and the *SlPSY1pro-*pAbAi did not grow, which confirmed the interaction of SlBZR1 and *SlPSY1* in Yeast (Figure 5C).

We further performed the dual luciferase reporter assays to test whether SlBZR1 could activate the transcription of *SlPSY1*. The promoter sequences of *SlPSY1* were fused successfully to the LUC reporter gene, co-transformed with an empty pGreenII 62-SK vector or pGreenII 62-SK-SlBZR1 vector into N. benthamiana leaf epidermal cells. The results showed that SlBZR1 significantly activated the promoter of the *SlPSY1* (1.55-fold) compared with the activity obtained with the control sample (Figure 5D). Thus, the results demonstrated that SlBZR1 positively regulates the expression of *SlPSY1* and carotenoid accumulation in ripening tomato fruits.

### SlBIN2 negatively regulates tomato fruit ripening

To further investigate the molecular mechanism of the SlBZR1-mediated regulation of tomato fruit ripening, a Y2H screen of tomato cDNA library was conducted to identify proteins interacting with SlBZR1. We successfully obtained several clones using Y2H screening, and one of them encoding a protein belonging to the Glycogen Synthase Kinase 3 (GSK3)-like kinases caught our attention. A BLAST search of the Phytozome database revealed that the GSK3-like kinase sequence was identical to *SlSK21/BIN2*, encode protein of 437 amino acids, with calculated molecular weights of 49.31 kDa. On the basis of a phylogenetic tree, the homologous genes of GSK3-like kinase family in Arabidopsis and tomato were divided into four subgroups, suggesting that *SlBIN2* was the homologous gene of *AtSK21/BIN2* in tomato, which is a negative regulator of BR signaling (Supplemental Figure S6, A and B). In order to confirm that SlBIN2 protein interacts with SlBZR1, we conducted a yeast two-hybrid assays on stringent selective medium (SD/-Trp-Leu-His-Ade) and X-α-gal. The result demonstrated that SlBIN2 interacted with SlBZR1, consistent with the interaction of BIN2 and BZR1 in Arabidopsis (Supplemental Figure S6C).

To gain insight into the biological function of SlBIN2 in tomato fruit ripening, we generated *SlBIN2*-OE transgenic lines and *SlBIN2* knock-out mutants using CRISPR/Cas9 system in AC. Three transgenic lines (OE-1, OE-2 and OE-4) with elevated *SlBIN2* gene expression were selected for further analysis (Supplemental Figure S6D). At the same time, two homozygous *bin2* mutants (*bin2-5* and *bin2-10*) with 10 bp and 2 bp deletions that led to translational frame shifts were obtained, which encode truncated SlBIN2 proteins with 33 and 10 aa, respectively (Supplemental Figure S6E). These results suggest that *bin2-5* and *bin2-10* are loss-of-function mutants suitable for the functional analysis of *SlBIN2*.

We observed and recorded the ripening characteristics of *SlBIN2*-OE and *bin2* mutant fruits. Compared with the WT plants, the fruit ripening progress was significantly inhibited in the *SlBIN2*-OE lines and promoted in the *bin2* mutant lines (Figure 6, A and B). *SlBIN2*-OE fruits required an additional 1-2 days, while *bin2* mutant fruits showed 1-2 days earlier from anthesis to the breaker stage when compared with WT fruits (Figure 6C). The peaks of climacteric ethylene production delayed about 1-2 days in *SlBIN2*-OE fruits and occurred 1-2 days earlier in *bin2* mutant fruits than in WT (Figure 6D).

**Figure 6.**
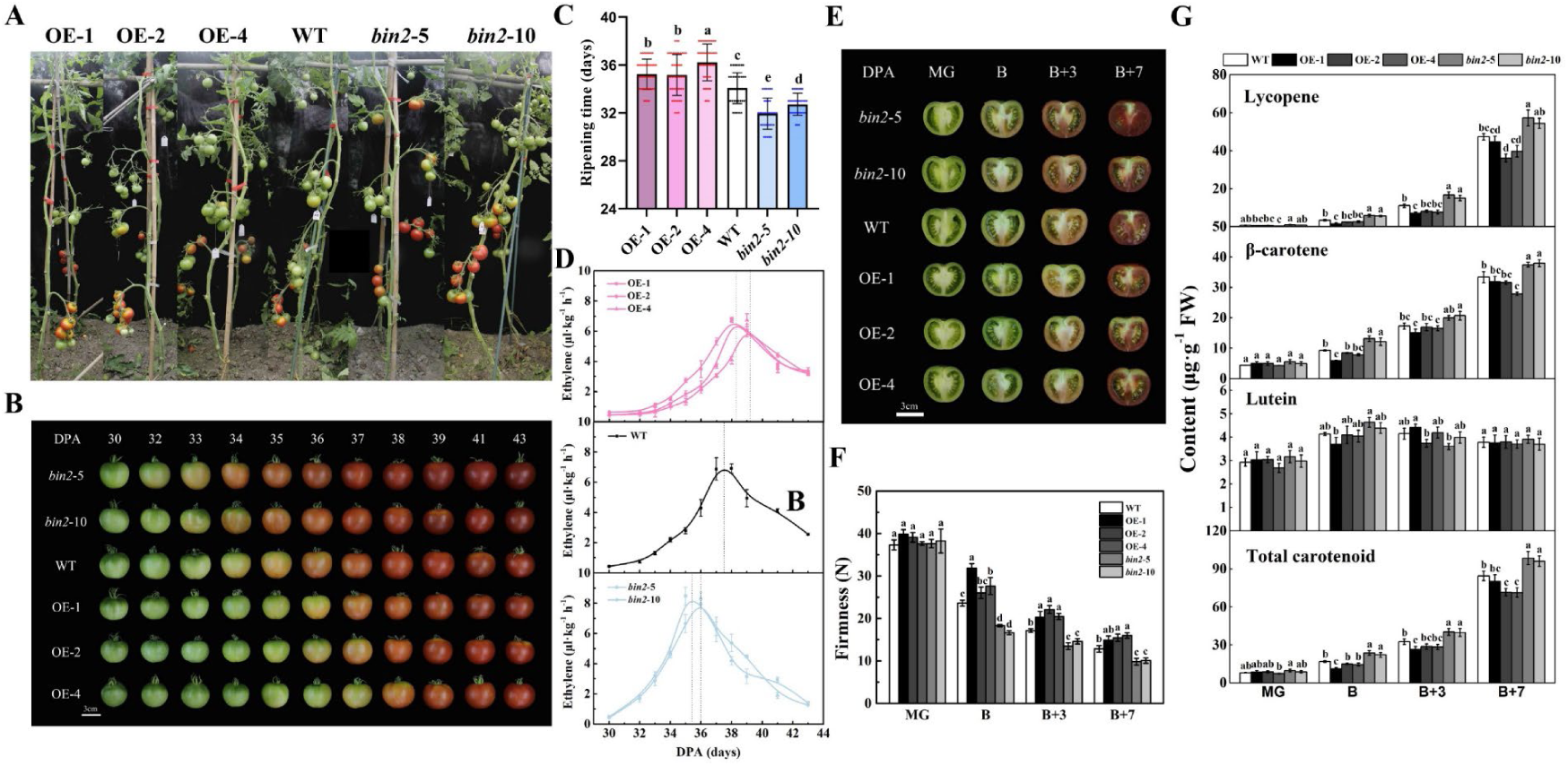
SlBIN2 delays tomato fruit ripening and reduces carotenoid accumulation. (A) Phenotypes of *SlBIN2*-OE and *bin2* mutant lines. (B) Ripening phenotype of *SlBIN2*-OE and *bin2* mutant fruits. Fruits from *SlBIN2*-OE lines show a delayed ripening phenotype while *bzr1* mutant lines ripen earlier. (C) Ripening time (days from anthesis to breaker) of WT, *SlBIN2*-OE and *bin2* mutant fruits. (D) Ethylene production of WT, *SlBIN2*-OE and *bin2* mutant fruits at different developmental stages. (E) Cross-sectioned WT, *SlBIN2*-OE and *bin2* mutant fruits at four different developmental stages. (F) Fruit firmness and carotenoids content (G) in WT, *SlBIN2*-OE and *bin2* mutant fruits at four different developmental stages. MG, mature green stage; B, breaker; B + 3, 3 d after breaker stage; B + 7, 7 d after breaker stage. Biological replicates (3-4 fruits per fruit ripening stage) were performed in triplicate, and the data are presented as means ± SE. The asterisks indicate statistically significant differences between the WT and transgenic fruits (P < 0.05).

To study the effects of *SlBIN2* on main physiological characteristics of tomato fruit ripening, we also measured fruit firmness and carotenoid content of *SlBIN2*-OE, *bin2* mutant and WT fruits. The results showed that the firmness of *SlBIN2*-OE fruits was significantly enhanced compared with that of WT fruit at the B, B+3 and B+7 stages, while the firmness of *bin2* mutant fruits was reduced than that of WT fruit at the same stage (Figure 6F). The contents of lycopene, β-carotene and total carotenoids in *SlBIN2*-OE fruits were lower than those in WT fruits, whereas *bin2* mutant fruits produced more carotenoids than WT fruits, especially lycopene (Figure 6G). Consistent with the observed delayed ripening and ethylene production as well as reduced carotenoid levels, the relative expression levels of the ethylene and carotenoid biosynthesis genes *SlACO1, SlACO3* and *SlPSY1* were lower in *SlBIN2*-OE fruits compared to WT fruits (Supplemental Figure S7 A, B and C). Taken together, these results indicated that SlBIN2, as a negative regulator of BR signaling, inhibits fruit ripening and carotenoids accumulation in tomato.

## Discussion

Brassinosteroids (BRs) are known as a group of growth-promoting steroid hormones which are required for the regulation of a variety of physiological processes during plant growth and development, including, seed germination, root growth, stem elongation, leaf morphogenesis, stomatal formation, and floral development [34]. Here, our study demonstrates that BR signaling is essential for regulating tomato fruit ripening by interaction with ethylene signaling pathway.

### Growth-promoting phytohormone BR is essential for tomato fruit ripening

Previous reports showed that BRs were involved in the growth and development of tomato fruit. Firstly, exogenous application of BRs to tomato plant enhanced the height of plants as well as the yield and quality of fruits. An increase in fruit number and the levels of lycopene and β-carotene were observed in tomato plants treated with 28-homobrassinolide [55, 56]. Moreover, tomato fruit ripening was significantly inhibited and promoted by Brz and EBR, respectively [41, 57]. Secondly, several BR-deficient and BR-insensitive mutants identified in tomato showed severe dwarfing, delayed fruit ripening and reduced fruit quality [58, 59]. d^x^, a tomato BR-deficient mutant with an impaired Dwarf gene encoding a BR C-6 oxidase, resulting in severe dwarfism. d^x^ fruits showed delayed ripening and reduced fruit yield and dry mass, which could be partially rescued by BR application [58, 60-62]. Thirdly, intense biosynthesis of BRs was detected during tomato fruit development, suggesting BRs was required for tomato fruit development [58, 63]. These studies illuminated that BRs play roles in tomato development, while the effect of endogenous BRs level on tomato fruit ripening is largely unknown.

Our previous results showed that the transcript levels of BR biosynthetic gene *SlDWARF* and *SlCYP90B3* were up-regulated, followed by an increase in the contents of endogenous BL and CS at the onset of tomato fruit ripening [48]. Additionally, overexpression of *SlCYP90B3* or *SlBRI1* enhanced endogenous BR levels or BR signaling, which promoted tomato fruit ripening [48, 50]. These results are consistent with the identification of several BR-deficient and BR-insensitive mutants showing delayed fruit ripening [58, 59, 62]. Moreover, our study reveals that overexpression of the positive regulator of BR signaling *SlBZR1* or knock-out of the negative regulator of BR signaling *SlBIN2* promote tomato fruit ripening (Figure 2, A and B; Figure 6, A and B). In contrast, *bzr1bes1* and *SlBIN2-OE* lines display delayed fruit ripening (Figure 1B; Figure 6, A and B). Our previous results indicate that SlBES1 promotes tomato fruit softening through transcriptional inhibition of *PMEU1* without affecting nutritional quality. The current research demonstrates that SlBZR1 functions as a master regulator of tomato fruit ripening, which directly binds and transactivates ethylene biosynthetic genes *SlACO1* and *SlACO3* and carotenoid biosynthetic genes *SlPSY1* to regulate ethylene production and carotenoid accumulation, thus promoting tomato fruit ripening. Moreover, our results reveal that SlBIN2 which interacts with SlBZR1 as a negative regulator of BR signaling, inhibits SlBZR1-mediated tomato fruit ripening and carotenoid accumulation. Together, these findings indicate that BR signaling promotes fruit ripening.

### Interplay of the BR and ethylene signaling pathway for regulation of tomato fruit ripening

In maturation process of climacteric fruits promoted by exogenous BR, the endogenous BR content did not change significantly in mango, banana and jujube [42, 44, 64], which increased in tomato and persimmon [43, 48] and decreased in apple and pear [53]. Similarly, the endogenous BR levels were found to be strongly elevated at the onset of fruit ripening in grape berries [46], implying that the mechanisms underlying BRs in controlling fruit ripening might be different depending on the fruit type.

As one of the primary initiators and regulators of maturation process in climacteric fruits, ethylene plays a key role in promoting ripeness and coloration of climacteric fruits, and its absence or decrease dramatically delays the ripening process [65]. Previous studies have demonstrated that BRs can directly regulate the ripening of non-climacteric fruits, but whether BRs regulate the ripening of climacteric fruits directly or via controlling ethylene biosynthesis remains unclear [66]. Interestingly, one study in Arabidopsis proposed that BRs at a low concentration negatively control ethylene production, while BRs at high concentrations positively regulate ethylene production [67, 68]. However, we cannot conclude that BRs either positively or negatively regulate ethylene biosynthesis in a dose-dependent manner, since the effects of of BRs upon different levels on ethylene biosynthesis are still absent in tomato fruits.

In this work, we found that exogenous application of eBL markedly promotes the production of ethylene, whereas PCZ treatment inhibits the climacteric peak of ethylene production (Supplemental Figure S1A). Furthermore, *SlCYP90B3*-OE accelerated tomato fruit ripening and advanced the onset of the climacteric peak of ethylene production, while *SlCYP90B3*-RNAi retarded tomato fruit ripening and caused a lower peak of ethylene production (Supplemental Figure S1C). Moreover, overexpression of either *SlDWARF, GhDWF4* or *SlBRI1* in tomato, results in elevated levels of ethylene production and earlier ripening as well as enhanced carotenoid accumulation [47, 49, 50]. In addition, *SlBZR1-OE* or *SlBIN2-KO* accelerated ethylene production and enhanced carotenoid accumulation with up-regulation of ethylene and carotenoid biosynthetic genes (Figure 2 and 6). In contrast, dual *SlBZR1/SlBES1*-KO or *SlBIN2-OE* tomato delayed ethylene production peaks and lower the content of carotenoids (Figure 1 and 6). These results suggest that BRs promote tomato fruit ripening possibly through activating the expression of ethylene biosynthetic genes and ultimately the ethylene production.

### SlBZR1 and SlBES1 function redundantly in tomato fruit ripening *via* distinct regulatory mechanisms

BR signaling is dependent on a series of phosphorylation events to modulate the function of BZR1/BES1 transcription factors that regulate the expression of BR-responsive genes [69, 70]. In this study, *SlBZR1* and *SlBES1*, the homologous gene of *AtBZR1* and *AtBES1*, were isolated from tomato. The expression of *SlBZR1* and *SlBES1* increased during the earlier stage of tomato fruit ripening, which suggests that *SlBZR1* and *SlBES1* are crucial regulators in the process of fruit ripening. Dual *SlBZR1/SlBES1*-KO tomato retarded fruit ripening with delayed ethylene production peaks and lower carotenoid content. Considering *SlBES1* promotes tomato fruit softening through transcriptional inhibition of *PMEU1* without affecting carotenoid levels and ethylene production, which indicates that *SlBZR1* might be essential in tomato fruit ripening and carotenoids accumulation [54].

Our study indicates that *SlBZR1-OE* promoted tomato fruit ripening with elevated ethylene and carotenoid contents, conversely, *SlBZR1-KO* retarded fruit ripening and ethylene production as well as lower carotenoid levels. Unraveling the molecular events underlying *SlBZR1* action by integrating RNA-seq and ChIP-seq data, revealed that knockout of *SlBZR1* results in massive transcriptomic reprogramming involving several families of transcription factors and a large number of fruits ripening related genes that are under the direct regulation of *SlBZR1*. The high number of DEGs involved in ethylene biosynthesis pathway and carotenoids metabolism pathway are influenced in the *SlBZR1*-KO lines support the notion that *SlBZR1* is a master regulator of tomato fruit ripening, in line with the increase in ethylene production and carotenoid accumulation.

This study showed that SlBZR1 positively regulated the expression of *SlACO1, SlACO3* and *SlPSY1* by directly binding to the E-boxes in the promoters of these genes during tomato fruit ripening (Figure 4 and 5). Similarly, in Arabidopsis roots, BES1/BZR2 increase ethylene production for gravitropism by directly binding to the E-boxes in the promoter of *ACO1* [71]. In contrast, BZR1 negatively regulates *ACO4* expression by binding to the BRRE-box of the *ACO4* promoter in arabidopsis seedlings [72]. Furthermore, BZR1 functions as a transcription activator and repressor on different promoters [38]. In this study, *SlBZR1* and *SlBES1*, as a transcriptional activator and repressor, promote tomato fruit ripening by increasing the biosynthesis of ethylene and carotenoid, and inhibiting pectin de-methylesterification, respectively. However, regarding BRs accelerate tomato fruit ripening [41, 48], we suggest that the transcriptional activator *SlBZR1* plays a major role in tomato fruit ripening.

We observed that SlBZR1 interacts with the SlBIN2 which functions as a negative regulator of BR signaling upstream. Previous studies have demonstrated that the presence of BR results in dephosphorylation of BZR1, and the dephosphorylated BZR1 then enters into the nucleus to regulate expression of its target genes [34, 73, 74]. The sub-cellular localization results showed that SlBZR1 are nucleus- and cytoplasm-localized proteins, and EBL treatment induced the nuclear localization of SlBZR1 proteins, whereas the interaction between SlBIN2 and SlBZR1 resulted in the phosphorylated SlBZR1 moving into cytoplasm, thus affecting SlBZR1 binding to the promoter of its target genes (Supplemental Figure S6D). These findings are similar to the results that BIN2-induced phosphorylation inactivates BZR1 by promoting their cytoplasmic retention via 14-3-3 proteins [74, 75]. Moreover, *SlBIN2-OE* delayed ethylene production peaks and lower the content of carotenoids, while *SlBIN2-KO* accelerated ethylene production and enhanced carotenoid accumulation (Figure 6). In combination with above observations, it can be concluded that SlBIN2induces the cytoplasmic retention of phosphorylated SlBZR1 via interaction with it, thereby inhibiting transcriptional activation of *SlACO1, SlACO3* and *SlPSY1* by SlBZR1.

In conclusion, our results support a model that BR signaling regulates tomato fruit ripening (Figure 7). When endogenous BRs are present at low levels, SlBIN2 is constitutively active and interacts with SlBZR1 to promote degradation of cytoplasmic located phosphorylated SlBZR1, thereby inhibit its DNA binding activities, thus retard tomato fruit ripening and decrease the levels of carotenoids. when endogenous BR levels are present at high levels, a series of phosphorylation and dephosphorylation reactions are occurred, leading to the inactivation and degradation of dephosphorylated SlBIN2. SlBZR1 is then rapidly dephosphorylated by BIN2 as substrate and moved into the nucleus to active the transcription of ethylene biosynthetic genes *SlACO1* and *SlACO3* and carotenoid biosynthetic gene *SlPSY1* by direct binding to their promoters, leading to a burst of ethylene production and carotenoids accumulation, thus promoting fruit ripening in tomato.

**Figure 7.**
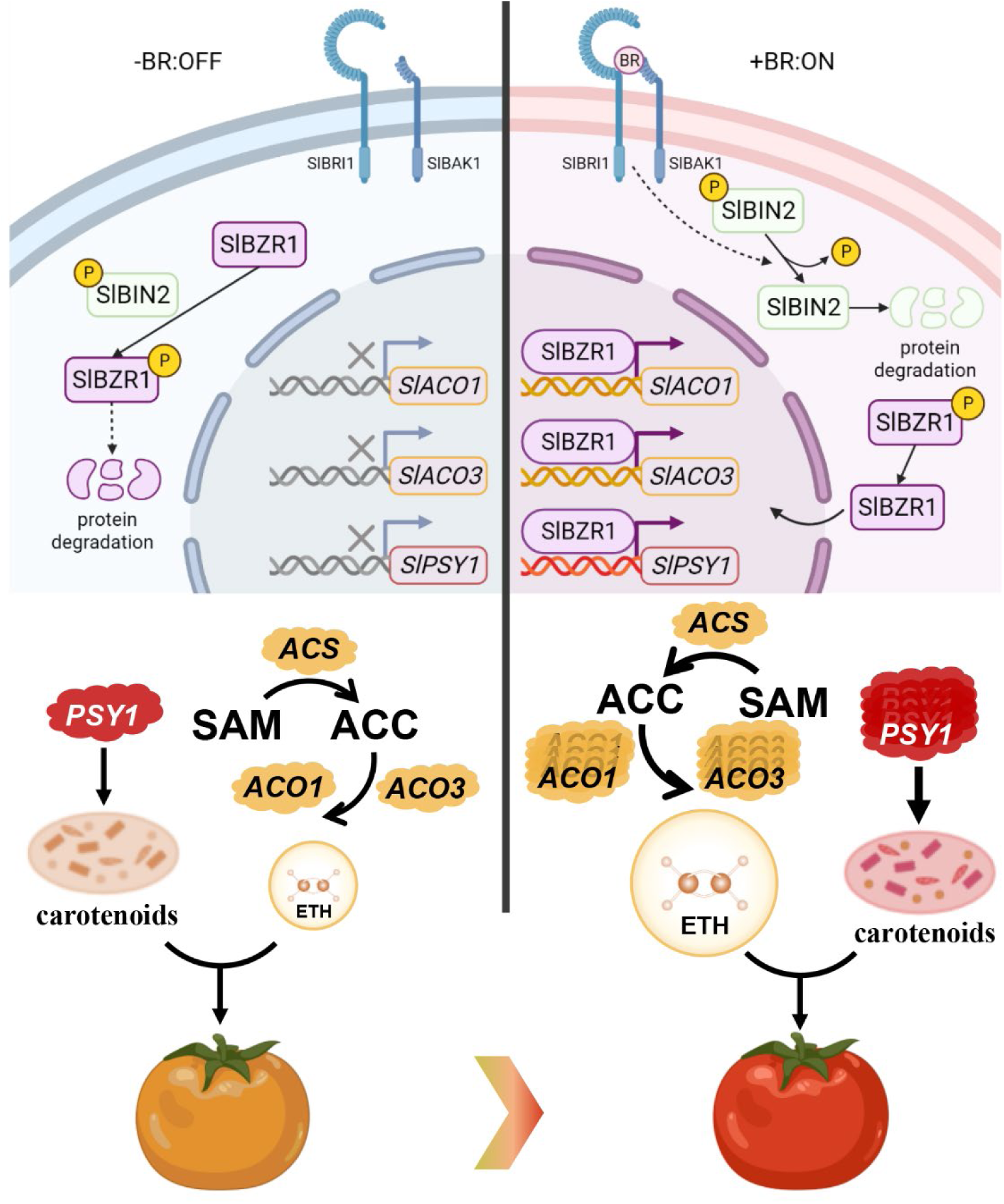
A proposed regulatory mechanism of BR on tomato fruit ripening and carotenoid accumulation. When BRs (brassinosteroids) are absent (left), BRI1 (Brassinosteroid-Insensitive1) and its co-receptor BAK1 (BRI1 Associated receptor Kinase 1) are kept inactive. BIN2 (Brassinosteroid-Insensitive 2) is constitutively active and phosphorylates BZR1 (Brassinazole-resistant1) transcription factor to promote its degradation and 14-3-3-mediated cytosolic retention, and to directly inhibit their DNA binding activities. When BRs are present (right) and bind to the extracellular domains of BRI1 and its co-receptor BAK1 to activate the two receptor kinases, a series of phosphorylation and dephosphorylation reactions are occurred, leading to the degradation of dephosphorylated SlBIN2. Phosphorylated SlBZR1 is dephosphorylated and moved into the nucleus to active the transcription of ethylene biosynthetic genes (*SlACO1* and *SlACO3*) and carotenoid biosynthetic gene *SlPSY1* by direct promoter binding, leading to a burst of ethylene production and carotenoids accumulation.

## ACKNOWLEDGEMENTS

We thank Tomato Genetics Resource Center (University of California, Davis, CA) for providing tomato seeds used in our study. We also thank Prof. Qi Xie (Chinese Academy of Sciences) and Prof. Yaoguang Liu (South China Agriculture University) for kindly providing vectors for *pYAO*:hSpCas9 and pYLCRISPR/Cas9 plasmids, respectively. This research was supported by National Natural Science Foundation of China (Key Program, No.31830078), the Ministry of Agriculture of China (2016ZX08009003-001), Zhejiang Provincial Ten-thousand Program for Leading Talents of Science and Technology Innovation (2018R52026) and Zhejiang Provincial Natural Science Foundation of China (No. LZ15C150001).

## AUTHOR CONTRIBUTION

Q.W., L.L., and F.M. designed the research. F.M., H.L., S.H., C.J., M.Z., S.L., Y.L., J.L., Y.J., M.W., Z.S., and Y.M. performed the research. F.M. H.L., and S.H. analyzed data. F.M., L.L., and Q.W. wrote the manuscript.

## DECLARATION OF INTERESTS

The authors declare no competing interests.

### Resource availability

#### Lead Contact

Further information and requests for resources and reagents should be directed to and will be fulfilled by Qiaomei Wang.

#### Materials Availability

This study did not generate new unique reagents.

#### Data and Code Availability

The RNA-seq data have been deposited at the NCBI Sequence Read Archive (SRA) database (http://www.ncbi.nlm.nih.gov/sra/) with the BioProject ID PRJNA635540 and are publicly available as of the date of publication. Accession numbers are listed in the key resources table. This paper does not report original code. Any additional information required to reanalyze the data reported in this paper is available from the lead contact upon request.

## EXPERIMENTAL MODEL AND SUBJECT DETAILS

### Plant Materials and Growth Conditions

We used *Solanum lycopersicum* cv. Ailsa Craig (AC) as the control genotype in the present study. The following tomato genotypes were used in this study: *CYP90B3-OE, CYP90B3-RNAi, SlBIN2*-OE, *bin2, SlBZR1*-OE, *bzr1 and bzr1bes1*. Transgenic plants *CYP90B3-OE* and *CYP90B3-RNAi* were generated during previous studies [48]. Plants were cultivated under a 16 h photoperiod (22/28°C, night/day). *Solanum lycopersicum* flowers were labeled at anthesis and the number of tagged fruits was limited to fewer than four per cluster. The fruits were collected after 30 to 46 DPA stage for the different fruit ripening stages. The fruit ripening stages, including mature green (MG) stage, breaker (B) stage, pink (P) stage, and red ripe (R) stage, which were defined based on fruit color as described previously [76]. At the MG stage, fruits reach full size and display jelly placental tissues. The fruits displaying their first signs of color change were identified as B stage, 3 and 7 days after B stage were classified as P and R stage, respectively. Three biological replicates for pericarp tissues of the fruits (each biological replicate consisting of at least three fruits) were collected after harvesting, frozen in liquid nitrogen immediately, and then stored at -80°C for further tests.

### Accession Numbers

The accession numbers for the genes described in this report are as follows: *SlBZR1* (Solyc12g089040), *SlBES1* (Solyc04g079980), *SlPSY1* (Solyc03g031860), *SlACO1* (Solyc07g049530), *SlACO3* (Solyc07g049550), *SlBIN2* (Solyc07g055200), *SlCYP90B3* (Solyc02g085360), *ACTIN7* (Solyc03g078400), *ACTIN2* (Solyc11g005330).

## METHOD DETAILS

### Vector Construction and Plant Transformation

To generate the *35S*_*pro*_:*SlBZR1*-eGFP construct, the full-length coding sequence (CDS) of *SlBZR1* without termination codon was amplified via PCR, cloned into pQB-V3, and recombined with the binary vector pK7FWG2.0 (*35S* promoter, N-eGFP). To generate the *35S*_*pro*_:*SlBIN2*-myc construct, *SlBIN2* CDS was PCR amplified, cloned into pQB-V3, and recombined with the binary vector PGWB17 (*35S* promoter, c-4myc). All vector constructs were generated following standard molecular biology protocols and using Gateway (Invitrogen) technology. Primers used for vector construction are listed in Supplemental Table. All above constructs were introduced into tomato cultivars AC via *Agrobacterium tumefaciens* LBA4404-mediated transformation. Transformants were selected based on their resistance to kanamycin [77]. Homozygous T2 or T3 transgenic plants were used for phenotypic and molecular characterization. T2 progeny of independent transgenic lines *SlBZR1*-OE (OE-8, OE-9 and OE-14) as well as *SlBIN2*-OE (OE-1, OE-2 and OE-4) were used in this study.

### Generation of *slbzr1, slbin2 and slbzr1 slbes1* Using CRISPR/Cas9 Technology

CRISPR-P (http://cbi.hzau.edu.cn/crispr/) was used to design the single guide RNA (sgRNA) sequence that target either *SlBZR1* or *SlBIN2* and two guide RNA sequences which targeted both *SlBZR1* and *SlBES1* genes. Sequences of the sgRNAs are listed in Supplemental Table. The CRISPR/Cas9 system was used to generate the *slbzr1* and *slbin2* mutant as previously described. The sgRNA expression cassette with target sequence were assembled to *pYAO*:hSpCas9 binary plasmid [78]. The CRISPR/Cas9 mutagenesis of *slbzr1 slbes1* was performed by expressing two guide RNA sequences which targeted both *SlBZR1* and *SlBES1* genes in pYLCRISPR/Cas9 vector [79]. After confirmation by sequencing, three final binary vectors containing sgRNA and Cas9 was transferred into *Agrobacterium tumefaciens* LBA4404, then introduced into AC as described previously. CRISPR/Cas9-induced mutations were genotyped by PCR amplification and DNA sequencing. The induced genomic mutations and the primers used for confirming mutation are given in Fig. 1 and Supplemental Table, respectively. Homozygous T2 transgenic plants *bzr1-5, bzr1-28, bzr1-29, bin2-5* and *bin2-10* as well as *bzr1bes1-1* and *bzr1bes1-2* were selected for further experiments.

### Chemical treatment

The fruits of AC and *SlBZR1-OE* (OE-14) were harvested at mature green stage (the MG stage). The fruits of AC were treated with 3 μM 24-epibrassinolide (EBL, Sigma, St. Louis, MO) or 5 μM Pcz as well as the combination of 3 μM eBL and 5 μM PCZ, respectively [51]. Fruits treated with solvent were used as a control. After the treatments, the fruits were stored at at 24 °C and 80% relative humidity and sampled at 1st, 3rd, 6th, and 9th day for further tests. For 1-MCP treatment, the fruits of AC and *SlBZR1-OE* was placed in 25 L containers and treated with 1 μL L^-1^ 1-MCP for 16 h. After treatment with 1-MCP, the fruits were removed from the containers and stored at 24 °C and 80% relative humidity and collected at 0, 1st, 4th, 7th, and 10th day for further tests.

### Measurement of ethylene production

Ethylene production of tomato fruit was detected according to former reports with some minor modifications. Five fruits were enclosed in 1.0 L airtight container at 20 °C for 2 h, 1 mL of the headspace gas was withdrawn from the container with a 1 mL syringe and immediately injected into a gas chromatograph (Shimadzu GC-17A, Kyoto, Japan), consisting of a flame ionization detector and a hp-5 column (50 m × 0.32 mm × 1.05 μm) with a detection temperature of 200 °C. The ethylene production was calculated according to commercial standards. Fresh tissues of fruit were mixed frozen in liquid nitrogen and stored at -80 °C until use.

### Firmness determination

The fruits of wild type and transgenic lines were harvested at each development stage. Firmness was measured at the fruit equatorial region with a TA-XT2i texture analyzer (Stable Micro Systems Ltd., Godalming, UK). The penetration depth was set at 10 mm and the penetration speed of a 7.5mm probe was set at 1 mm s^-1^. The unit of force for firmness is Newton.

### HPLC analysis of carotenoids

The extract and analysis of carotenoids was conducted as early described using high performance liquid chromatography (Shimadzu, Kyoto, Japan) with minor modification [30]. About 0.5 g of tomato fruit powder was extracted with 30 mL of hexane, acetone and ethanol (1:1:1, v/v/v). The mixture was shaken and then 15 mL of double distilled water added. The extracts were centrifuged at 12000 g for 15 min at 4 °C. The supernatant was eluted through a 0.22 μm filter membrane then evaporated to dryness with nitrogen blowing instrument. The condensed extract was dissolved in tetrahydrofuran: acetonitrile: methanol (15:30:55, v/v/v) with 1.5 mL dissolution. Then the solutions were loaded into liquid phase bottle for HPLC analysis.

The analysis of carotenoids was determined using HPLC instrument (Shimadzu, Kyoto, Japan), consisting of a SPD-M20A diode array detector and chromatographic column C18 (5 μm particle size, 4.6 mm × 250 mm, Elite analytical instruments Co., Ltd., Dalian, China), The mobile phase was methanol: acetonitrile (90:10, v/v) containing 0.05 % TEA. The current velocity and the wavelength were set as 1.2 mL/min and 475 nm, respectively. Peak area of carotenoids was selected based on the spectral characteristics and then calculated according to standard compounds (lutein, lycopene, and β-carotene; Sigma, USA).

### RNA extraction and qRT-PCR analysis

For RNA extraction, 0.1 g of leaves or fruits was mixed with 1 mL RNAiso plus according to manufacturer’s instruction (Takara, Kusatsu, Japan), then RNA was reverse-transcribed into cDNA using PrimeScript RT reagent with gDNA Eraser (Takara, Kusatsu, Japan). TB Green (Takara, Kusatsu, Japan) was then used in Step One Real-Time PCR System (Applied biosystem, CA, USA) for relative quantitative PCR (qPCR). The gene-specific primers used were listed in Supplemental Table.

### Subcellular localization

The *35S*_*pro*_:*SlBZR1*-eGFP and *35S*_*pro*_:*SlBIN2*-myc constructs was co-infiltrated with a mCherry-labeled nuclear marker NF-YA4-mCherry into *N. benthamiana* leaves using *A. tumefaciens*-mediated infiltration [80]. The N. benthamiana plants were kept in the dark for 48 h after infiltration. Then EBR (10 μM) was injected into the infiltrated *N. benthamiana* leaves and imaging was performed 12 h after EBR treatment. GFP fluorescence was observed under a confocal microscope (TCS SP8; Leica Wetzlar, Germany). For green fluorescence observation, the excitation wavelength was 488 nm and the emission wavelengths were 520∼540 nm; for red fluorescence observation, the excitation wavelength was 561 nm and the emission wavelengths were 610∼630 nm. *35S*_*pro*_:eGFP was used as a control. All transient expression assays were repeated at least 3 times, and the representative results were shown.

### RNA-seq

Total RNA was extracted from tomato fruits of WT and *bzr1* at MG and B stages and Illumina MiSeq library was constructed as manufacturer’s instructions (Illumina, San Diego, CA, USA) described. Then each sample library was sequenced with the Illumina Miseq platform (Shanghai Personal Biotechnology Cp., Ltd., Shanghai, China).

### ChIP-seq and ChIP-qPCR Assays

Chromatin immunoprecipitation sequencing was performed as previously reported [81]. In brief, tomato fruits of *SlBZR1-*OE at MG stage were collected from at least six different plants, sliced into small pieces and fixed in 1x PBS with 1% formaldehyde for 15 min under vacuum and ground to fine power under liquid nitrogen. ChIP assay was performed in three independent biological replicates starting with 2 g of each sample tissue. Nuclei were purified using a one-step Percoll gradient centrifugation as previously described. ChIP-Seq was performed with the anti-GFP antibody (Millipore) using transgenic plants expressing their cDNA fused to GFP under the 35S promoter. For conventional TruSeq libraries, the reverse crosslinked DNA was end-repaired, dA-tailed, ligated to Y-adapters, and PCR amplified prior to sequencing. For the more recent Tn5-based ChIP-Seq libraries, 1-2 uL of purified and assembled transposome (1 OD) was added to the washed magnetic beads in the presence of 30 uL 1x Tn5 buffer (10 mM MgCl_2_, 25 mM Tris pH 8.0, 10% DMF). The tagmentation reaction was performed at 37 °C for 30 min. The beads were then washed with low salt, high salt, and TE washing buffers and reverse crosslinked overnight. The purified DNA was then PCR amplified using N70x and N50x index primers. Two to three biological replicates were sequenced for each experiment.

For ChIP-qPCR, 3 g of 0.5 cm^2^ fruit pieces of *SlBZR1-*OE were collected at mature green stage and cross-linked using 1%(v/v) formaldehyde under vacuum for 10 min and ground to powder in liquid nitrogen. Then, the chromatin complexes were isolated following former research, sonicated with Biorupter plus (Diagenode, Belgium, Antwerp), and immunoprecipitated with 5 μg Mouse monoclonal anti-GFP antibody (clone 9E10, IgG, Roche, Switzerland). About 200∼300 bp of ChIP DNA and input DNA were recovered and dissolved in water for further qPCR analysis [82]. The primers used for ChIP-qPCR were designed around E-box in the enriched regions and are listed in Supplemental Data. Each ChIP value was normalized to its respective input DNA value. The fold enrichment on each candidate genes promoter was calculated against the ACTIN7 or ACTIN2, respectively. Mean value of two technical replicates was recorded for each biological replicate. The values of three independent biological replicates were collected and error bars represented the standard error from these experiments.

### Dual luciferase transactivation assay

The whole length of SlBZR1 CDS was fused with pGreen II-SK vector to generate an effector construct. The promoter was cloned into the vector pGreen II-LUC to generate a reporter construct using gene-specific primers. These constructs were introduced into Agrobacterium cell GV3101 together with the pSoup helper plasmid. Tobacco leaves were agroinfiltrated with the indicated plasmids, and harvested 3 d later. NightOWL II LB983 Ultrasens backlit (Berthold Technologies, Bad Wildbad, Germany) with Indigo software was used to capture LUC expression image and to quantify LUC luminescence intensity. 100 mM luciferin was sprayed on leaves and incubated in dark for 5 min before detection. Six independent determinations were performed.

### Yeast one-hybrid assay

The SlBZR1 CDS was amplified and inserted into pGADT7 to generate a prey construct. The promoter fragments of *SlPSY1* (1,835-bp upstream of the predicted translation start site), *SlACO1* (546-bp upstream of the predicted translation start site) or *SlACO3* (2828-bp upstream of the predicted translation start site) were cloned separately into the pAbAi vector to create bait constructs. After digesting with BstB1, these bait constructs were used to transform yeast strain Y1HGold and integrated into the yeast genome to create the bait reporter strains. The minimal inhibitory concentration of aureobasidin A (AbA) was screened to avoid self-activation. The prey vector and the empty vector (AD), serving as the negative control, were transformed separately into the bait reporter strains and grown on SD medium lacking Leu (SD/-Leu) plates with or without AbA at 30°C for 72 h. The positive clones were picked and diluted in double-distilled water to a final concentration of OD600 = 1.0. The suspension was spotted on SD/-Leu medium with or without AbA to assess protein-DNA interactions.

### Yeast two-hybrid assay

The TAD deletion constructs of SlBZR1 were used for the Y2H screening of the pGADT7-based tomato cDNA library, which was generated with mRNAs isolated from toamto leaves, roots, stems, flowers and fruits. The SlBZR1 CDS was introduced into the pGBKT7 vector and then used as bait to screen the cDNA library based on the Matchmaker Gold Yeast Two-Hybrid System (Clontech). Yeast transformants were exhaustively selected on SD/-Ade/-His/-Leu/-Trp/X-α-Gal medium. Putative SlBZR1 interacting clones were characterized and sequenced.

Full length coding sequence (CDS) of SlBIN2 was ligated to the binding domain (BD) in the pGBKT7 vector, and full-length CDS of SlBZR1 was introduced into the activation domain (AD) vector (pGADT7). The BD and AD vectors were co-transformed into the Y2H Gold yeast strain (AH109). Transformed yeast cells were grown on SD/-Leu/-Trp medium and SD/-Ade/-His/-Leu/-Trp medium containing x-a-gal to determine protein-protein interactions.

### Statistical analysis

The experimental data were analyzed with SPSS 19.0 software. Multiple comparisons were subjected to ANOVA using Duncan’s test, while pairwise comparisons were computed using Student’s *t*-test (*P* < 0.05). Statistically signifificant differences (*P* < 0.05) are represented by different lowercase letters.

## Notes

### Competing Interest Statement

The authors have declared no competing interest.

### Summary of Updates

Supplemental files updated

