## Supplemental figures for "Brassinosteroids signaling component SlBZR1 promotes fruit ripening in tomato"

### Supplementary Figure S1-S7

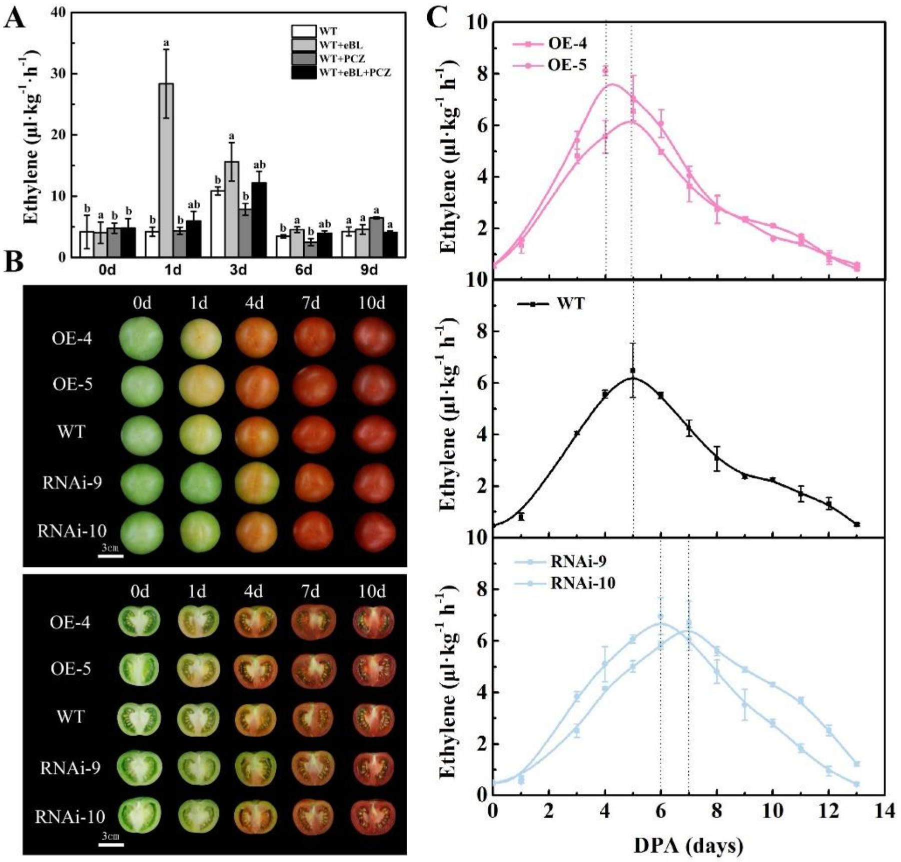


**Supplemental Figure S1. Ethylene production in tomato fruits treated with either individual or combination of 24-epibrassinolide (EBL) and propiconazole (PCZ), as well as *CYP90B3*-OE, WT (wild type) and *CYP90B3*-RNAi fruits. (Supports Figure 1)**

(A) Tomato fruits were collected at mature green stage (30 DPA, days post anthesis), ethylene production of WT fruits treated with EBL and PCZ alone or combination, as well as *CYP90B3*-OE, WT and *CYP90B3*-RNAi fruits (B, C) stored for different days at room temperature were measured. 0 d indicates the fruit harvested on the day of mature green stage. Three biological replicates were analyzed as described in the Methods section. Values represent means ± SD. Statistical significance was determined using a Student's $t$-test ($P<0.05$).

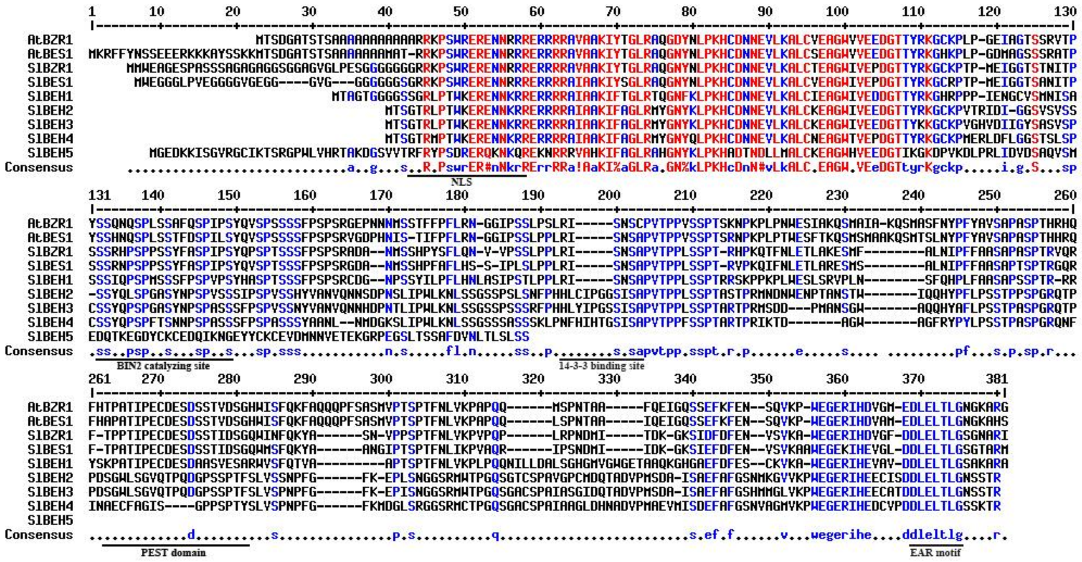


**Supplemental Figure S2. Amino acid sequence alignment of SlBZR1/SlBES1 with homologs from Arabidopsis and tomato. (Supports Figure 1)**

Multiple sequence alignment of SlBZR1/SlBES1 and related proteins from Arabidopsis and tomato. The horizontal lines mark the four conserved domains. The important structures for BZR proteins including the NLS (nuclear localization signal domain), BIN2 catalyzing site and PEST domain are represented with black lines.

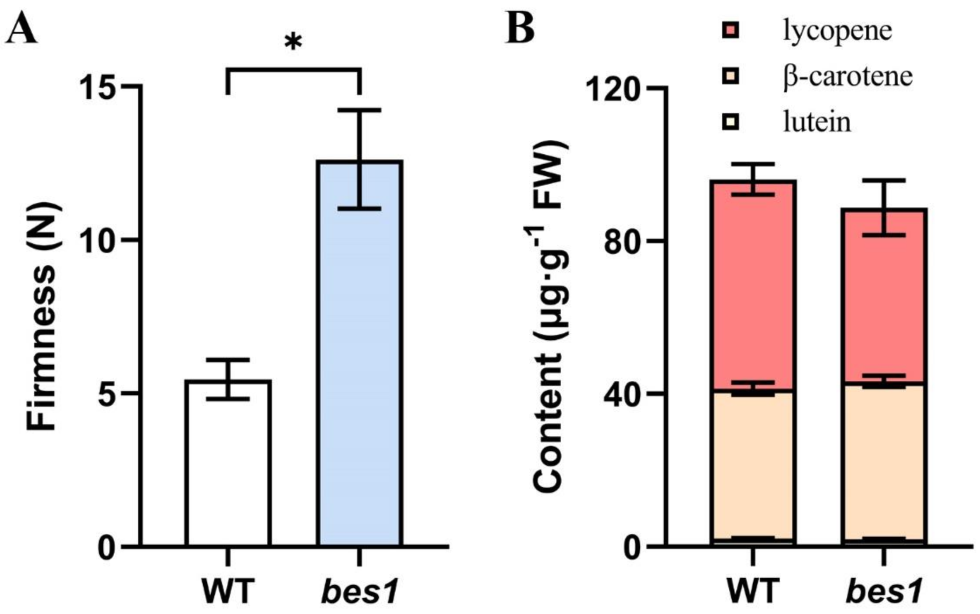


**Supplemental Figure S3. *SlBES1* promotes tomato fruit softening without affecting carotenoids accumulation and ethylene production. (Supports Figure 2 and Figure 3)**

(A) Fruit firmness and carotenoids content (B) in WT and *bes1* fruits at R stage.

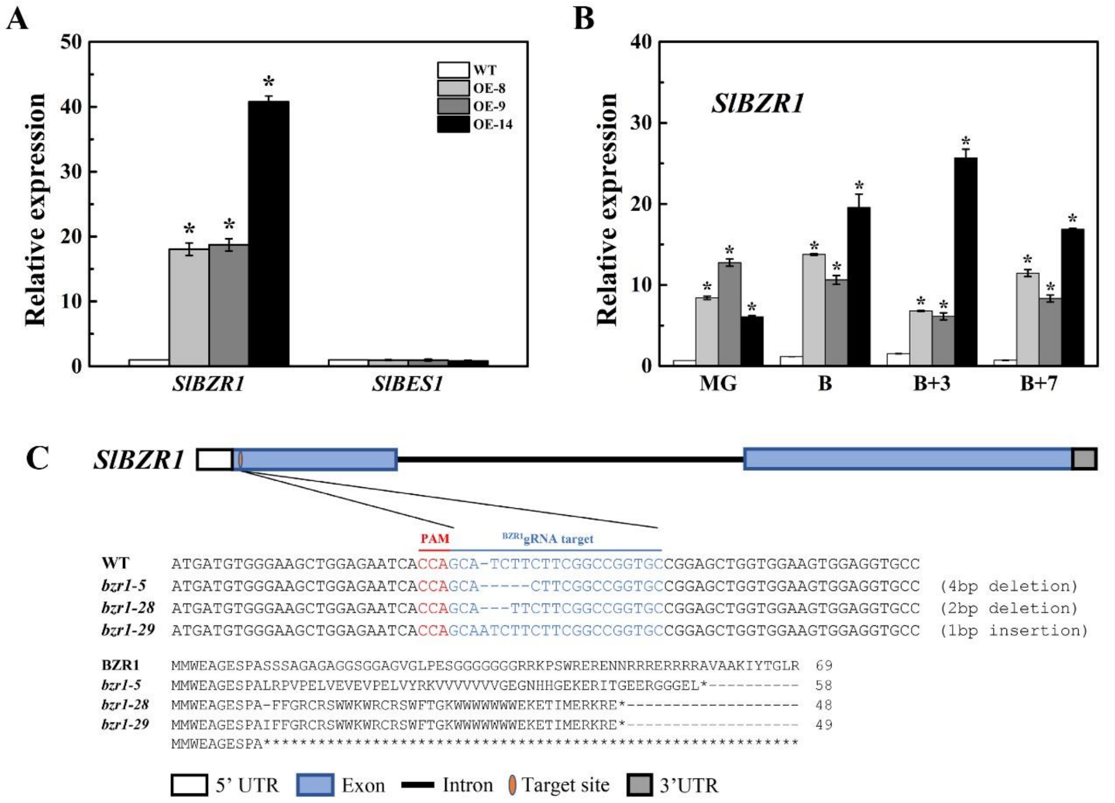


**Supplemental Figure S4. The generation of SlBZR1-OE transgenic lines and CRISPR/Cas9-engineered mutations in SlBZR1. (Supports Figure 2 and Figure 3)**

(A) Expression levels of *SlBZR1* and *SlBES1* in the leaves of *SlBZR1*-OE lines. (A) Expression levels of *SlBZR1* in the fruits at different developmental stages of *SlBZR1*-OE lines. (C) Genotype of mutations in the SlBZR1 locus generated by the CRISPR/Cas9 genome editing system. One target sequence was designed to specifically target the first exon. The red and blue letters indicate the sequences of protospacer adjacent motif (PAM) and target, respectively. The mutations in the transgenic plants were confirmed by sequencing the genomic regions flanking the target sites. The sequences of WT plants and three homozygous mutant lines *bzr1-5*, *bzr1-28* and *bzr1-29* are shown. Asterisks indicate statistical significance using Student's t-test ($P < 0.05$).

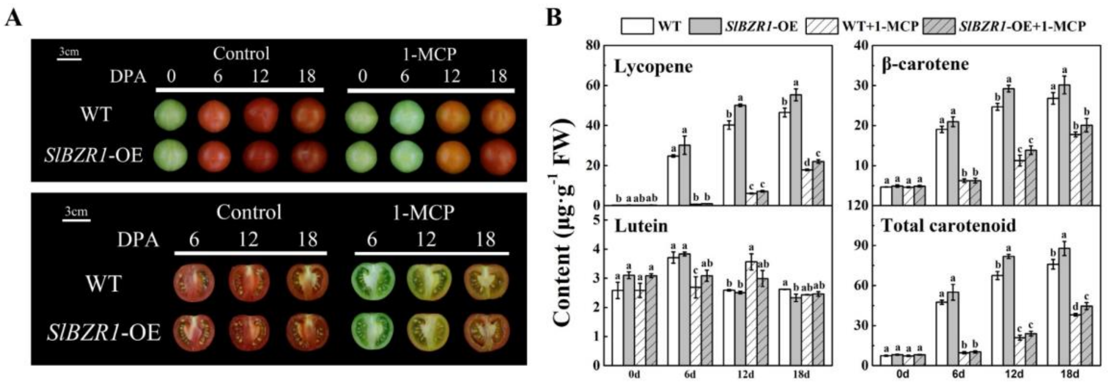


**Supplemental Figure S5. SlBZR1 promotes carotenoid accumulation in a partially ethylene-dependent manner. (Supports Figure 4 and Figure 5)**

(A) Tomato fruits were collected at MG stage (30 DPA, days post anthesis), visual appearance of wild-type and SlBZR1-OE fruits treated with or without 1-MCP stored for different days at room temperature. (B) Carotenoid accumulation in WT and *SlBZR1*-OE fruits with or without 1-MCP treatment. Here, 1 $\mu L \cdot L^{-1}$ 1-MCP was applied as treatment. Three biological replicates (n = 3) were used for each analysis ($P < 0.05$; Student's t-test). The error bars indicated the standard deviations.

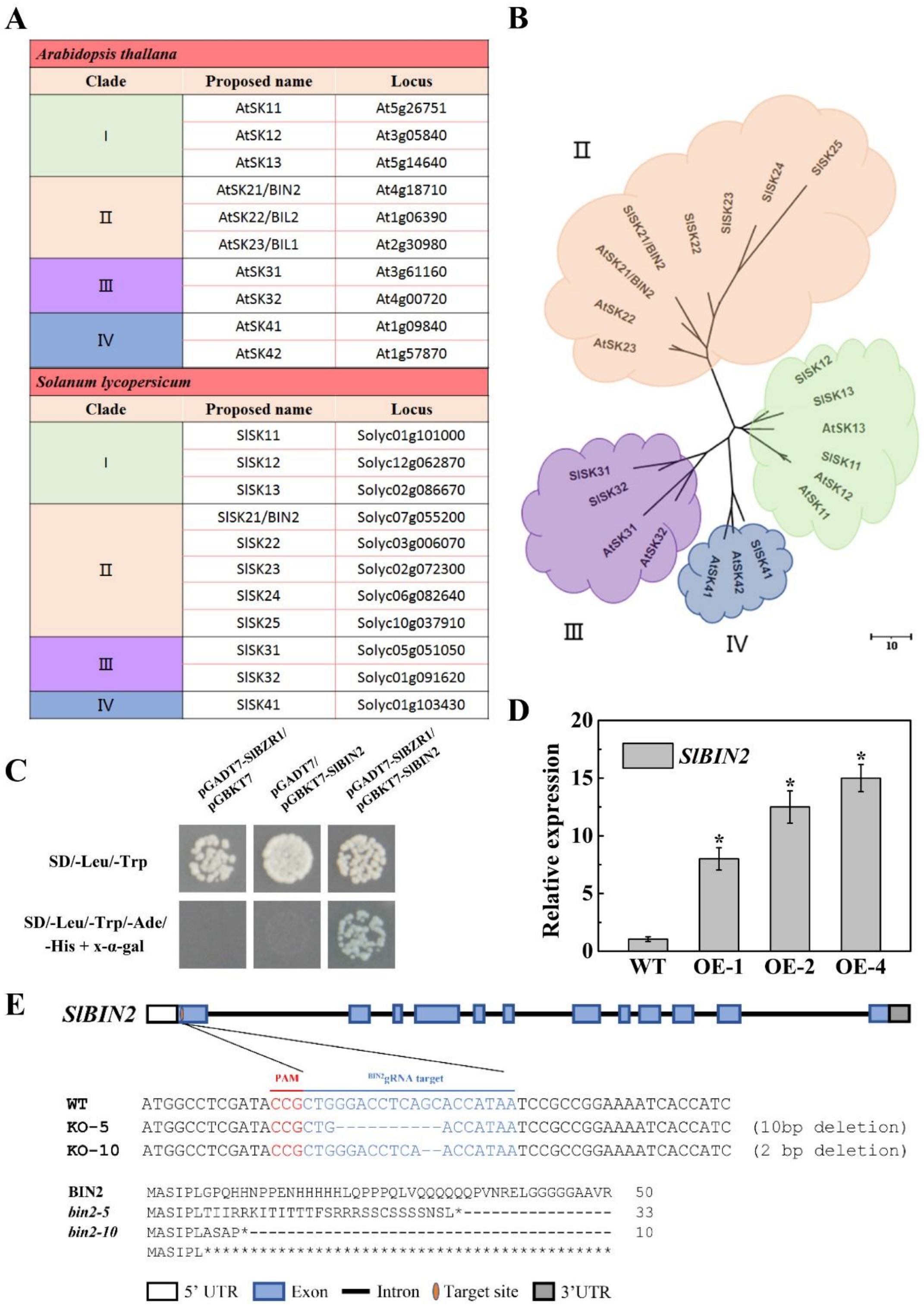


**Supplemental Figure S6. Tomato GSK3-like Kinases SlBIN2 Interacts with SlBZR1. (Supports Figure 6)**

(A) The GSK3-like kinase family of Arabidopsis thaliana and tomato is classified into four subgroups (I–IV). We propose to unify their official name as AtSKs (Arabidopsis thaliana Shaggy/GSK3-like kinases) or SlSKs (tomato Shaggy/GSK3-like kinases). (B) Phylogenetic analysis of Arabidopsis and tomato GSK3-like kinases. Amino acid sequences of GSK3-like kinases, which were obtained from TAIR and Sol Genomics Network resources, were aligned with ClustalW. The evolutionary history was inferred using the Maximum Parsimony method (Nei and Kumar, 2000). Evolutionary analyses were conducted in MEGA6 (Tamura et al., 2013). The scale bar indicates 10 amino acid substitutions. (C) Yeast two-hybrid (Y2H) assay for the interaction between SlBIN2 and SlBZR1. Full length coding sequence (CDS) of SlBIN2 was fused with the BD in pGBKT7, and full-length CDS of SlBZR1 was fused with the AD in pGADT7. Transformed yeast cells were grown on SD/-Leu/-Trp medium and SD/-Ade/-His/-Leu/-Trp medium containing x-a-gal to determine protein-protein interactions. (D) Gene expression levels of *SlBZR1* in leaves of *SlBIN2*-OE lines. (E) Genotype of mutations in the *SlBIN2* locus generated by the CRISPR/Cas9 genome editing system. One target sequence was designed to specifically target the first exon. The red and blue letters indicate the sequences of protospacer adjacent motif (PAM) and target, respectively. The mutations in the transgenic plants were confirmed by sequencing the genomic regions flanking the target sites. The sequences of the WT plants and two homozygous mutant lines (*bin2-5* and *bin2-10*) are shown.

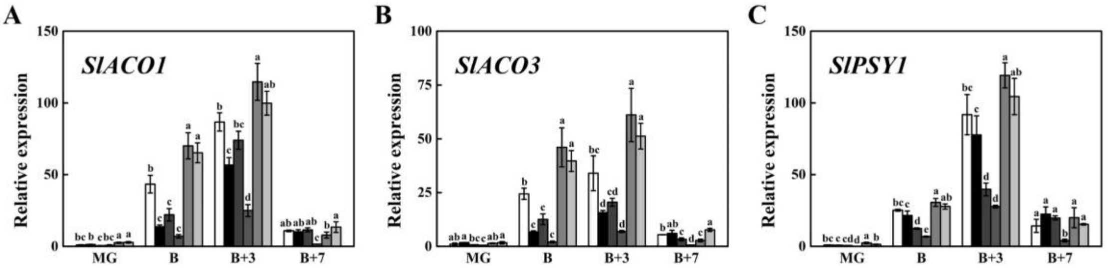


**Supplemental Figure S7. Expression levels of ethylene and carotenoid biosynthetic genes in *SlBIN2* overexpression and knockout tomato fruits. (Supports Figure 6)**

(A) Relative expression levels of *SlACO1*, *SlACO3* (B) and *SlPSY1* (C) in *SlBZR1*-OE and *bzr1* mutant fruits at different development stages. MG, mature green stage; B, breaker; B + 3, 3 d after breaker stage; B + 7, 7 d after breaker stage. Biological replicates (3-4 fruits per fruit ripening stage) were performed in triplicate, and the data are presented as means ± SE. The asterisks indicate statistically significant differences between the WT and transgenic fruits ($P < 0.05$).
